# MYC drives endothelial transformation and dedifferentiation in angiosarcoma

**DOI:** 10.64898/2026.10.07.757149

**Authors:** Chen Khuan Wong, Dan Li, Tomer Itkin, Cindy J. Lee, Nicholas A. Teri, Jiasheng Zhou, Dana M. Schoeps, Nikita R. Staniloae, Mohini R. Pachai, Makhzuna N. Khudoynazarova, Priya Pancholi, Woo Hyun Cho, Gabriella Bayshtok, Kae Kristoff, Selina Fan, Richard Lin, Shahin Rafii, Benedikt Bosbach, Scott W. Lowe, Cristina R. Antonescu, Ping Chi, Yu Chen

**Affiliations:** Human Oncology and Pathogenesis Program, Department of Medicine, Memorial Sloan Kettering Cancer Center, New York, NY 10065, USA; Division of Regenerative Medicine, Hartman Institute for Therapeutic Organ Regeneration, Ansary Stem Cell Institute, Department of Medicine, Weill Cornell Medicine, New York, NY 10065, USA; Department of Cancer Biology and Genetics, Memorial Sloan Kettering Cancer Center, New York, NY 10065, USA; Howard Hughes Medical Institute, Chevy Chase, MD 20815, USA; Department of Pathology, Memorial Sloan Kettering Cancer Center, New York, NY 10065, USA; Department of Medicine, Memorial Sloan Kettering Cancer Center, New York, NY 10065, USA; Department of Medicine, Weill Cornell Medical College, New York, NY 10065, USA

## Abstract

*MYC* amplification is the most common genetic alteration in radiation-associated angiosarcoma and is also present in a subset of *de novo* primary angiosarcoma, yet its oncogenic role in this vascular malignancy remains unclear. Here, we report that inducible, endothelial-specific *MYC* overexpression alone was sufficient to drive angiosarcomagenesis in mice. The resulting pulmonary angiosarcoma tumors exhibited impaired endothelial differentiation, indicative of MYC-driven lineage plasticity. Notably, *MYC* inactivation in dedifferentiated allografts induced growth arrest and restored endothelial gene expression, demonstrating MYC-dependent reversible cellular plasticity. Integrative analyses of MYC ChIP-seq, ATAC-seq, and RNA-seq revealed that overexpressed MYC preferentially occupied highly accessible promoters and activated genes involved in cell growth. Consistent with the murine findings, *MYC*-amplified human angiosarcomas were enriched for MYC target and cell-cycle programs and depleted of endothelial lineage signatures. *MYC* overexpression in human endothelial and angiosarcoma cells recapitulated these transcriptional changes, enhanced proliferation, and impaired endothelial tube formation. Together, our results establish MYC as an oncoprotein that promotes endothelial lineage plasticity to drive angiosarcoma initiation and progression.

## INTRODUCTION

Angiosarcoma (AS) is a rare, aggressive malignancy of endothelial origin that can arise in virtually any anatomical site [1, 2]. Patients with AS have a poor prognosis, with a 5-year overall survival rate of less than 30% [3]. Although most cases arise sporadically, therapeutic radiation is a well-established risk factor for secondary AS. A defining molecular feature of radiation-associated AS is focal amplification and overexpression of *MYC* [4, 5]. *MYC* amplification is also detected in approximately 13% of primary *de novo* AS cases [6]. Despite the prevalence of *MYC* amplification in AS, experimental models for defining the functional role of MYC overexpression in this disease have been lacking.

MYC is a basic helix-loop-helix leucine zipper transcription factor that regulates stem cell function and becomes oncogenic when aberrantly expressed in multiple cancer contexts [7]. Transgenic mouse models have demonstrated that *MYC* overexpression can drive malignancies in several tissues, including blood, liver, and prostate. Although MYC dysregulation is a defining oncogenic event in Burkitt lymphoma, MYC-induced tumorigenesis in many other tissues exhibits prolonged latency or requires cooperating oncogenic alterations [7]. Analysis of the MSK-IMPACT AS cohort showed that *MYC* amplification was associated with a lower tumor mutational burden and fraction of the genome altered [6]. These observations raised the possibility that endothelial cells are particularly susceptible to MYC-driven transformation and that *MYC* overexpression may require relatively few additional cooperating events to initiate AS. We therefore investigated whether inducible, endothelial-specific *MYC* overexpression was sufficient to drive angiosarcomagenesis in murine models.

## RESULTS

### *MYC* overexpression drives aggressive endothelial tumorigenesis *in vivo*

Vascular endothelial cadherin (VE-Cad), encoded by *Cdh5,* is expressed predominantly in endothelial cells. The *VE-Cad::CreER^T2^*allele has been widely used to achieve temporally controlled, endothelial-specific recombination [8, 9]. To induce human *MYC* expression reversibly in endothelial cells, we generated a genetically engineered mouse model (GEMM) combining 3 alleles: 1) *VE-Cad::CreER^T2^*, 2) *CAG-LSL*-*rtTA3-IRES-mKate2* [10] and 3) *TetO*- *MYC* [11] (**Fig. 1A**). We intraperitoneally administered tamoxifen in 6 to 8-week-old mice, an age where VE-Cad::CreER^T2^ expression is highly specific to endothelial cells. To identify cells that underwent Cre-mediated excision of the LSL cassette and expressed mKATE2, we analyzed dissociated lung cells by fluorescence-activated cell sorting (FACS) using the endothelial cell-specific *Pecam1*-encoded surface marker CD31 and mKATE2 fluorescence. mKATE2 expression was restricted to the CD31^+^ compartment, and approximately 20% of CD31^+^ cells were mKATE2^+^ (**Fig. 1B** & **supp. Fig. 1A**), demonstrating endothelial-specific recombination with modest efficiency.

**Figure 1:**
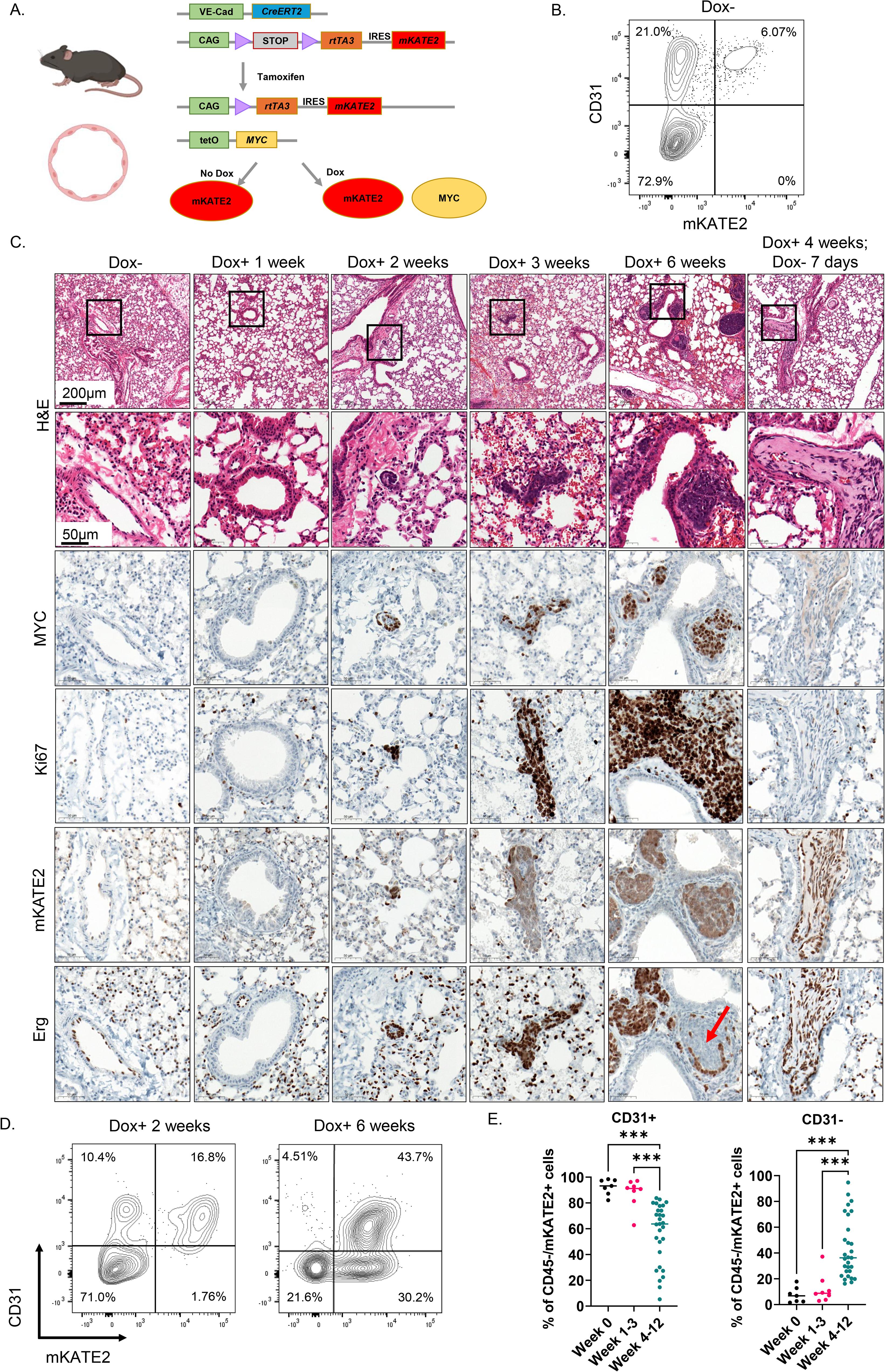
MYC overexpression transforms endothelial cells into angiosarcoma. **(A)** Schematic of the genetically engineered mouse model. *VE-Cadherin*-expressing endothelial cells were targeted for the tamoxifen-inducible expression of *rtTA3* and *mKATE2* and doxycycline-inducible expression of *MYC*. **(B)** Representative FACS analysis of CD31 expression and mKATE2 fluorescence among dissociated CD45^-^ lung cells from a tamoxifen- treated, doxycycline-naive mouse. **(C)** H&E and IHC staining of lung tissues from tamoxifen- treated mice after the indicated durations of doxycycline administration. The arrow indicates Erg^-^ tumor cells. **(D)** Representative FACS analysis of CD31 expression and mKATE2 fluorescence among dissociated CD45^-^ lung cells from tamoxifen-treated mice after the indicated durations of doxycycline administration. **(E)** Quantification of the CD31^+^ and CD31^-^ fractions among CD45^-^/mKATE2^+^ lungs cells after the indicated durations of doxycycline administration. Week 0, n=7; weeks 1-3, n=8; weeks 4-12, n=29. ***adjusted p < 0.001 (One-way ANOVA).

To activate *MYC* expression in mKATE2^+^ endothelial cells, mice were administered doxycycline in their drinking water and analyzed temporally after treatment initiation. Beginning two weeks after doxycycline administration, hematoxylin and eosin (H&E) staining of lung tissue revealed small clusters of neoplastic cells associated with vascular structures. These cells expressed mKATE2, MYC, and the endothelial-lineage transcription factor Erg (**Fig. 1C**). The neoplastic cells exhibited enlarged nuclei, prominent nucleoli, and a high Ki67 proliferation index, and the tumor nodules increased in size over time. By week 6, subsets of tumor cells had lost Erg staining (**Fig. 1C, arrow**). Erg^+^ and Erg^-^ tumor cells exhibited similar histological features and were found adjacent to one other within individual tumor nodules. Following doxycycline withdrawal, the tumor cells lost MYC and Ki67 expression and exhibited increased endothelial differentiation (**Fig 1C**).

To characterize changes in endothelial differentiation over time, we analyzed CD31 expression and mKATE2 fluorescence by FACS. In the absence of doxycycline, nearly all mKATE2^+^ cells were CD31^+^. Following doxycycline treatment, an mKATE2^+^/CD31^-^ population emerged and increased over time (**Fig. 1D and E**). The rapid development of these tumors demonstrates that *MYC* overexpression in pulmonary endothelial cells is sufficient to initiate angiosarcomagenesis, whereas the progressive emergence of mKATE2^+^/CD31^-^ and Erg^-^ tumor cells indicates loss of endothelial differentiation with sustained *MYC* overexpression. In addition to pulmonary angiosarcomas, tumors with high Ki67 expression and heterogenous Erg staining developed in the hearts of transgenic mice (**supp. Fig. 1B**).

### MYC induces transcriptional reprogramming of endothelial cells

To characterize the endothelial cell-state dynamics associated with *MYC* activation, we performed single-nucleus (sn) Multiome sequencing of endothelial lineage cells isolated from the lungs of transgenic mice (n=3) after 4 weeks of doxycycline treatment. We used FACS to remove dead cells and non-endothelial cells (CD31^-^/mKATE2^-^). The retained populations captured by FACS comprised endothelial cells that did not undergo Cre-mediated activation of rtTA3-IRES-mKATE2 (CD31^+^/mKATE2^-^), differentiated AS cells (CD31^+^/mKATE2^+^), and dedifferentiated AS cells (CD31^-^/mKATE2^+^).

We visualized the transcriptomes of the 4,866 nuclei on uniform manifold approximation and projection (UMAP) and identified eight transcriptional clusters by Leiden clustering, including four endothelial subclusters (**supp. Fig. 2A**). Based on endothelial marker and the *MYC* transgene expression, we consolidated these clusters into three major transcriptional cell states: normal endothelial cells (Normal), differentiated tumor (DiffT), and dedifferentiated tumor (DediffT) (**Fig. 2A-B and supp. Fig 2B-C**). *MYC*-negative Normal cells exhibited the highest expression of endothelial markers, including *Cdh5*, *Pecam1*, and *Erg*, whereas *MYC*-positive DiffT cells retained intermediate expression of these genes (**Fig. 2C**). In contrast, DediffT cells largely lost endothelial gene expression while exhibiting the highest levels of *MYC*, the proliferative marker *Mki67*, and cell cycle genes (**Fig. 2D**). Compared with Normal cells, DiffT and DediffT cells also exhibited higher RNA counts and greater number of genes detected per nucleus, consistent with prior reports of increased cellular RNA output in MYC-overexpressing cells **(supp. Fig. 2D**) [12, 13].

**Figure 2:**
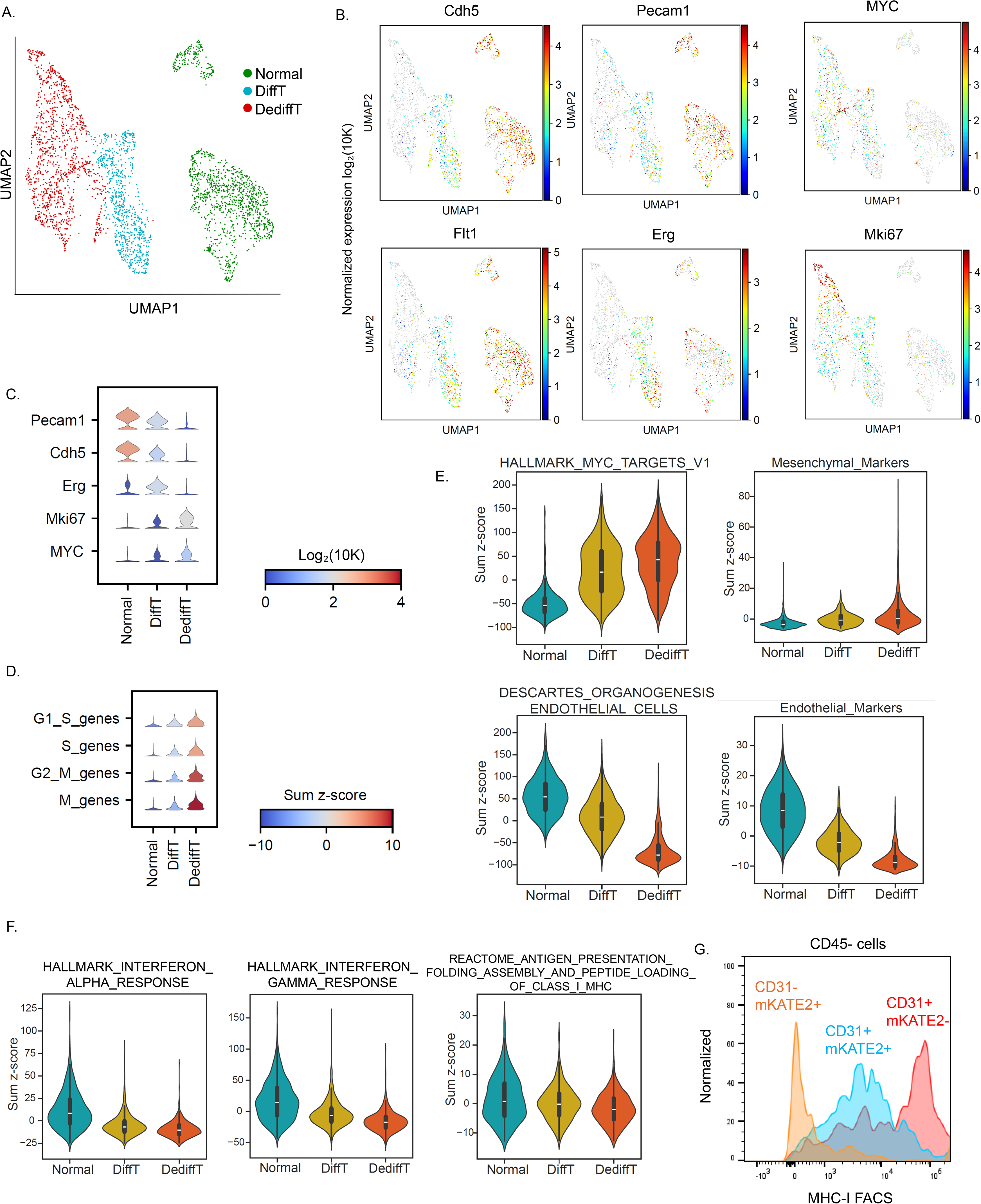
MYC induces transcriptional reprogramming of endothelial cells. **(A)** UMAP of single-nucleus RNA expression profiles from 4,866 nuclei obtained by Multiome sequencing of FACS-sorted CD31^+^/mKATE2^-^, CD31^+^/mKATE2^+^, and CD31^-^/mKATE2^+^ populations isolated from CD45^-^ lung cells pooled from n=3 mice. Nuclei are colored according to their assignment as normal endothelial (Normal), differentiated tumor (DiffT), or dedifferentiated tumor (DediffT) cells. **(B)** snRNA-seq UMAPs colored by expression of the indicated endothelial marker genes, *MYC*, and *Mki67*. **(C)** Violin plots showing normalized expression of the indicated endothelial marker genes, *Mki67*, and *MYC* in Normal, DiffT and DediffT cells. **(D)** Violin plots showing the sum z-scores for cell-cycle genes in the three cell states**. (E)** Violin plots showing sum z-scores for the indicated gene sets across the three cell states. **(F)** Sum z-scores for the indicated immune-related gene sets in the three cell states. **(G)** Representative FACS analysis of MHC-I expression in CD31^+^/mKATE2^-^, CD31^+^/mKATE2^+^, and CD31^-^/mKATE2^+^ cells among dissociated CD45^-^ lung cells from a mouse treated with tamoxifen and doxycycline for 6 weeks.

We next scored two Hallmark MYC target signatures and two endothelial cell-specific signatures. MYC target gene signatures were elevated in the neoplastic DiffT and DediffT cells, whereas endothelial lineage signatures progressively decreased across the Normal, DiffT, and DediffT states, suggesting that *MYC* activation is associated with loss of endothelial identity (**Fig. 2E**). We further annotated cell states using CellTypist with Tabula Muris as a reference [14, 15]. As expected, Normal and DiffT were predominantly classified as endothelial, whereas DediffT cells were assigned to multiple cell types, with the largest fractions classified as mesenchymal and endothelial cells (**supp. Fig. 2E**). Because endothelial cells can acquire mesenchymal features through endothelial-mesenchymal transition (EndoMT) during development, tissue repair, and disease pathogenesis [12], we examined the expression of mesenchymal markers, including *Acta2*, *Col1a1*, *Tagln*, and *Fn1*. Their expression was higher in DediffT than in DiffT and higher in DiffT than in Normal cells, indicating progressive acquisition of an EndoMT-like transcriptional program during MYC-driven transformation (**supp. Fig. 2F**).

In addition to losing endothelial lineage signatures, DiffT and DediffT cells exhibited reduced interferon-response and inflammatory gene-set scores, suggesting that MYC-driven transformation is associated with suppression of immune-related signaling programs **(Fig. 2F)**. MYC overexpression has been associated with downregulation of the major histocompatibility complex class I (MHC-1) expression and immune evasion in multiple cancer types [16–18]. Consistent with this, interferon alpha, interferon gamma, and class I MHC gene signatures were downregulated in DediffT and DiffT cells compared to Normal cells (**Fig. 2F**). We next measured cell surface MHC-I expression in dissociated lung cells by flow cytometry. CD31^+^/mKATE2^-^ Normal endothelial cells exhibited high MHC-I expression, whereas CD31^+^/mKATE2^+^ DiffT cells showed reduced expression. CD31^-^/mKATE2^+^ DediffT cells exhibited a further reduction in MHC-I expression, suggesting that *MYC* activation and endothelial dedifferentiation are associated with diminished antigen presentation capacity (**Fig. 2G**). To orthogonally validate the snRNA-seq findings, we performed bulk RNA-seq on FACS sorted normal endothelial cells (CD31^+^/mKATE2^-^), differentiated tumor cells (CD31^+^/mKATE2^+^), and dedifferentiated tumor cells (CD31^-^/mKATE2^+^). We observed concordant changes to those observed in snRNA-seq (**supp. Fig. 2G-I**).

### MYC-driven cell states exhibit distinct chromatin accessibility landscapes

To characterize chromatin accessibility following endothelial *MYC* overexpression, we analyzed the snATAC-seq data from the endothelial lineage cells isolated from the lungs. Graph-based clustering and UMAP visualization identified three major clusters (**supp. Fig. 3A**). Fragment-size distributions, transcription start site enrichment, and the number of unique fragments per nucleus were assessed as measures of data quality (**supp. Fig. 3B and 3C**). Integration of snRNA-seq expression profiles with the snATAC-seq data enabled annotation of these clusters as normal endothelial (Normal), differentiated tumor (DiffT), and dedifferentiated tumor (DediffT) cell states (**Fig. 3A**).

**Figure 3:**
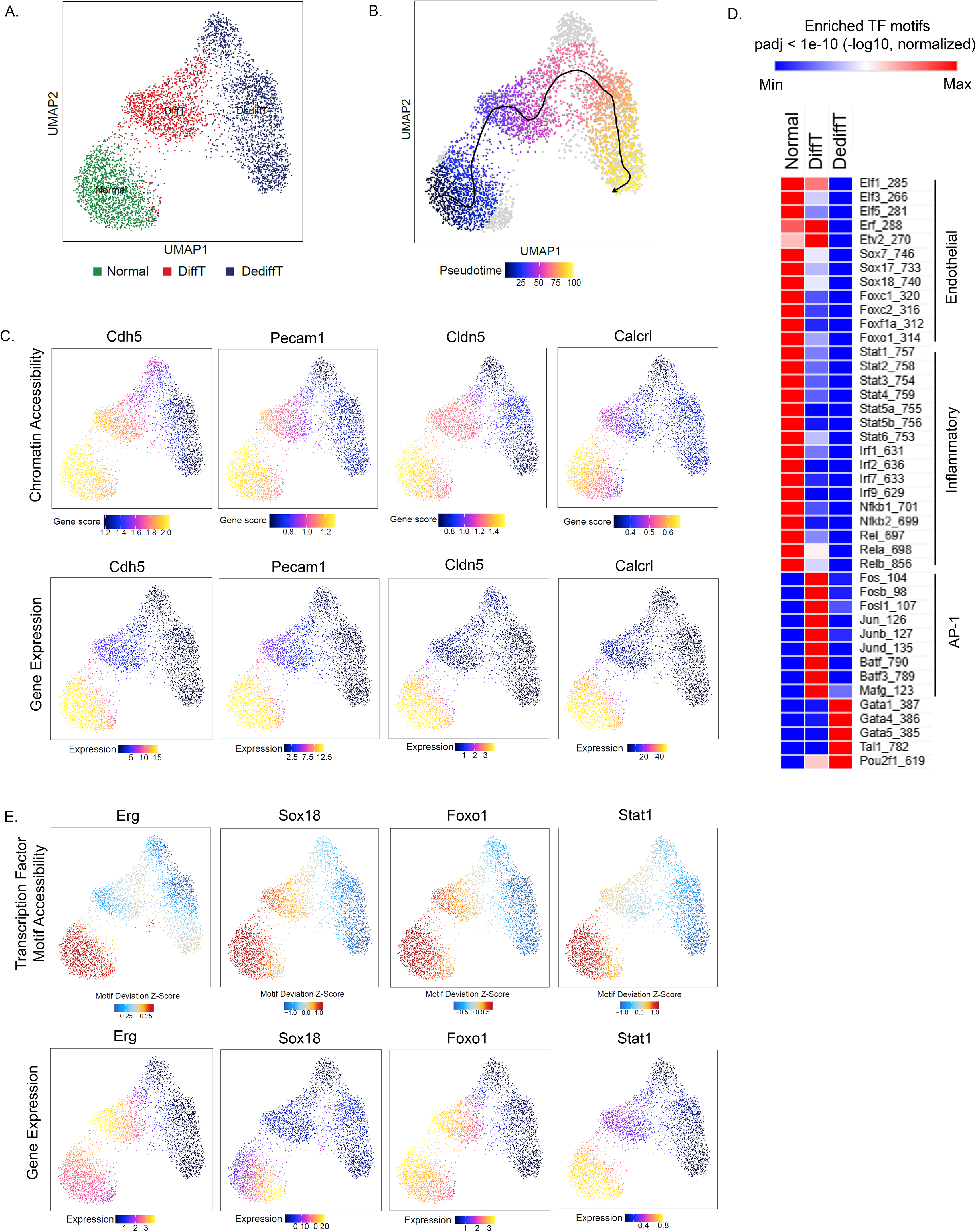
MYC overexpression alters the chromatin accessibility landscape that defines the endothelial lineage. **(A)** snATAC-seq UMAP of FACSorted endothelial-lineage cells isolated from the lungs of mice (n=3) showing three major clusters. **(B)** Integration of snRNA-seq data with the snATAC-seq embedding identified Normal, DiffT, and DediffT cell states. **(C)** Inferred trajectory from normal endothelial cells towards tumor dedifferentiation, displayed on the snATAC-seq UMAP and colored by pseudotime. **(D)** UMAPs showing gene-associated chromatin accessibility (top; GeneScoreMatrix, log₂-normalized counts) and integrated gene expression (bottom; GeneIntegrationMatrix normalized counts) for the indicated endothelial marker genes. **(E)** Heatmap of selected transcription factor motifs enrichments across the three cell states. Only motifs significantly enriched in at least one cell state (adjusted p < 0.05) are shown. Values represent −log10 (adjusted p-values), and colors are scaled relative to the minimum and maximum values for each motif. **(F)** UMAPs showing transcription factor motif accessibility (top; chromVAR motif deviation z-score) and integrated expression of the corresponding transcription factors (bottom; GeneIntegrationMatrix normalized counts) for factors implicated in maintaining endothelial identity.

We next performed pseudotime trajectory analysis, specifying the Normal state as the trajectory origin. The inferred trajectory progressed through the DiffT state and terminated at the DediffT state, consistent with progressive dedifferentiation during MYC-driven endothelial transformation (**Fig. 3B**). Normal endothelial cells exhibited high gene-associated chromatin accessibility at pan-endothelial markers, including *Cdh5, Pecam1, Cldn5,* and *Calcrl.* Accessibility at these loci progressively decreased in the DiffT and DediffT states, concomitant with reduced expression of the corresponding genes (**Fig. 3C**).

Motif enrichment analysis of cell-state-specific differentially accessible peaks revealed progressive depletion of motifs recognized by members of the Ets, Sox, and Fox transcription factor (TF) families, which contribute to the maintenance of the endothelial identity (**Fig. 3D**) [19, 20]. Motif associated with immune-regulatory transcription factors, including Stat and Irf family members, showed a similar pattern. Conversely, AP-1 family motifs were strongly enriched in the DiffT state, whereas Gata and Pou family motifs were enriched in the DediffT state.

To examine these regulatory programs at single-cell resolution, we projected chromVAR motif deviation z-scores and integrated expression of the corresponding transcription factors onto the snATAC-seq UMAP (**Fig. 3E and supp. Fig. 3D**). Motif accessibility and expression of TFs involved in endothelial development and identity, including Erg, Sox18, and Foxo1, were highest in the Normal state and progressively decreased across the DiffT and DediffT states. Motif accessibility and expression of immune-regulatory factors, including Stat1 and Irf1, were also reduced in tumor cells, consistent with attenuation of immune-related regulatory programs during dedifferentiation. Overall, MYC-driven endothelial transformation was accompanied by progressive loss of the chromatin accessibility landscape defining endothelial identity and acquisition of distinct tumor cell-state-specific regulatory programs.

### MYC is required to sustain angiosarcoma tumor growth

To determine whether MYC is required for tumor maintenance, we transplanted CD31^-^/mKATE2^+^ AS cells subcutaneously into syngeneic immunocompetent mice. After tumors were established, mice were randomized to continued doxycycline treatment or doxycycline withdrawal. Following doxycycline withdrawal and *MYC* inactivation, allografts exhibited reduced growth by day 2 and began to regress by day 4, demonstrating that established tumors are dependent on sustained *MYC* overexpression (**Fig. 4A**).

**Figure 4:**
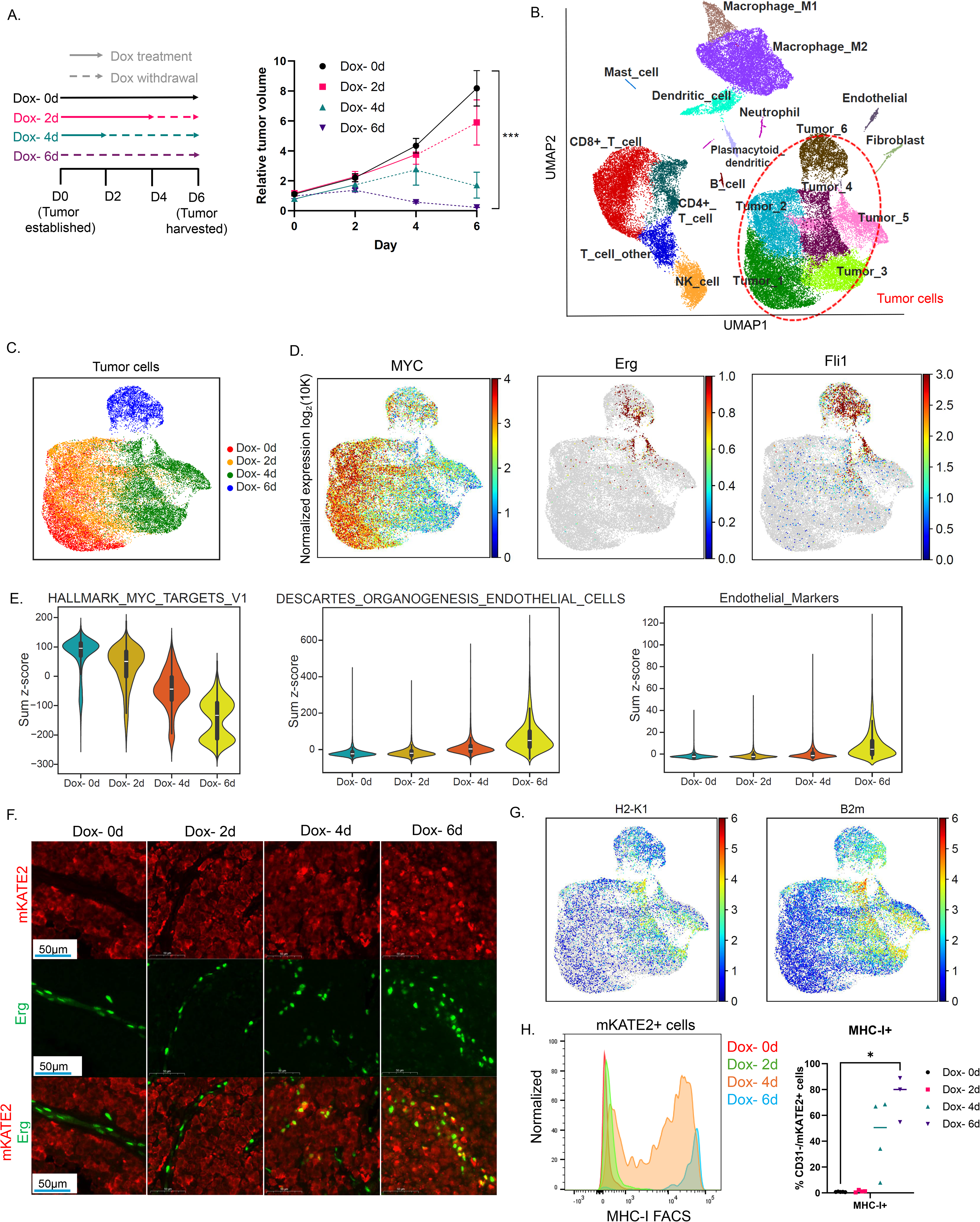
Response to MYC withdrawal in murine angiosarcoma. **(A)** Growth of established AS allografts in syngeneic immunocompetent mice maintained on doxycycline (Dox^+^, solid lines) or following doxycycline withdrawal (Dox^-^, dashed lines). Mean ± SEM, n=4 mice per group. *** p < 0.001 at day 6, two-tailed t-test. **(B)** UMAP of cells analyzed by scRNA-seq from AS allograftes collected from mice at baseline (n=3 mice) or 2 (n=2), 4 (n=2), or 6 days (n=3) after doxycycline withdrawal. Cells are colored by cluster identity. Tumor cells formed 6 clusters and aggregated closely (bottom right, outlined in red) while non-tumor cells formed distinct clusters. **(C)** UMAP of the tumor-cell compartment colored by the duration of doxycycline withdrawal. **(D)** UMAPs of the tumor-cell compartment colored by the expression of *MYC* and endothelial marker genes *Erg* and *Fli1*. **(E)** Violin plots showing sum z-scores for the indicated gene signatures in tumor cells at the specified times after doxycycline withdrawal. **(F)** Multiplex immunofluorescence staining of Erg and mKATE2 in AS allografts at the indicated times after doxycycline withdrawal. **(G)** UMAPs of the tumor-cell compartment colored by the expression of the antigen presentation genes *H2-K1* and *B2m*. **(H)** Flow cytometry analysis of cell-surface MHC-I expression on mKATE2^+^ tumor cells from allografts collected at the indicated times following doxycycline withdrawal. *p < 0.05 (Day 0 vs day 6, two-tailed unpaired t-test).

To characterize changes in tumor cell state following *MYC* inactivation, we performed single-cell (sc) RNA-seq on allografts collected at baseline or 2, 4, and 6 days after doxycycline withdrawal (**Fig. 4B & supp. Fig. 4A**). Because mKATE2^+^ tumor cells were scarce after 6 days of *MYC* inactivation, mKATE2^+^ tumor cells and mKATE2^-^ non-tumor cells were isolated by FACS and combined in equal numbers before scRNA-seq. UMAP analysis distinguished mKATE2^+^ tumor cells from the nonmalignant cells within the allografts (**Fig 4B**, **supp fig. 4B- C**). We first focused on the tumor cells that were separated into 6 transcriptional states. We found that cells at baseline predominantly enriched in clusters 1 and 2; day 2 cells enriched in clusters 2 and 4; day 4 cells enriched in clusters 3 and 5; and most day 6 cells enriched in a distinct cluster 6. We therefore examined temporal transcriptional changes according to the duration of *MYC* inactivation (**Fig. 4C**).

As expected, doxycycline withdrawal led to progressive downregulation of the *MYC* transcripts in tumor cells (**Fig. 4D**). After 6 days of *MYC* inactivation, the remaining tumor cells exhibited increased expression of endothelial markers, including *Erg* and *Fli1*, suggesting that the dedifferentiated tumor cells retain endothelial lineage memory and the capacity to reacquire features of endothelial differentiation. Similarly, we found upregulation of endothelial signatures and depletion of Hallmark MYC target and cell-cycle signatures following *MYC* inactivation (**Fig. 4E and supp. Fig. 4D**). Immunohistochemical analysis confirmed depletion of MYC and Ki67 proteins following doxycycline withdrawal, accompanied by increased Erg staining (**supp. Fig. 4E**). Multiplex immunofluorescence further demonstrated Erg expression in subsets of mKATE2^+^ tumor cells following doxycycline withdrawal (**Fig. 4F and supp. Fig. 4F**). Together, these results demonstrate that AS tumors depend on sustained *MYC* overexpression for growth and that *MYC* inactivation permits partial restoration of endothelial lineage identity.

We next investigated whether *MYC* inactivation altered tumor-intrinsic immune signaling and in the immune composition of the AS microenvironment. Following doxycycline withdrawal, tumor cells exhibited increased expression of inflammatory and interferon signaling signatures (**supp. Fig. 4G**), including upregulation of MHC class I genes H2-K1 and B2m (**Fig. 4G**). Flow cytometry confirmed progressive upregulation of cell-surface MHC-I on allografted tumor cells following *MYC* inactivation (**Fig. 4H**). Analysis of the immune compartment revealed increased proportions of CD4^+^ and CD8^+^ T cells and a decreased proportion of macrophages after 6 days of doxycycline withdrawal (**supp. Fig. 4H**). Flow cytometry and immunohistochemical staining further confirmed increased infiltration of CD8^+^ T cells in the allografts after 6 days of doxycycline withdrawal (**supp. Fig. 4I-J**).

Transcriptomic analysis of CD8^+^ T cells in *MYC*-active tumors revealed co-expression of cytotoxic granules, including *Nkg7* and *Gzmb,* and exhaustion markers, including *Pdcd1*, *Tox*, and *Havcr2*, consistent with a chronically stimulated, exhaustion-like state (**supp. Fig. 4K**). Following *MYC* inactivation, intratumoral CD8^+^ T cells exhibited increased expression of effector molecules, including *Ifng*, *Prf1*, and *Ccl3*, with cytotoxicity scores peaking at day 4 (**supp. Fig. 4L**). Together with increased interferon-response programs and MHC class I expression in tumor cells, these findings suggest that *MYC* inactivation enhances tumor immunogenicity and promotes a more cytotoxic CD8^+^ T-cell state.

### MYC binds accessible promoters and activates canonical growth programs in angiosarcoma

To define the cell-autonomous mechanisms by which MYC regulates gene expression, we derived a cell line, referred to here as MYC-AS, from allografts established using CD31-/mKATE2+ FACS-sorted cells. MYC-AS cells were maintained in doxycycline-supplemented medium to sustain *MYC* expression. MYC protein was depleted within 24h of doxycycline withdrawal, resulting in reduced cell growth *in vitro* (**Fig. 5A-B**). Bulk RNA-seq comparing *MYC*-active (Dox^+^) cells and *MYC*-inactive cells 24h after doxycycline withdrawal (Dox^-^) revealed depletion of Hallmark MYC target signatures and enrichment of endothelial and immune-related signatures following *MYC* inactivation (**Fig. 5C**).

**Figure 5:**
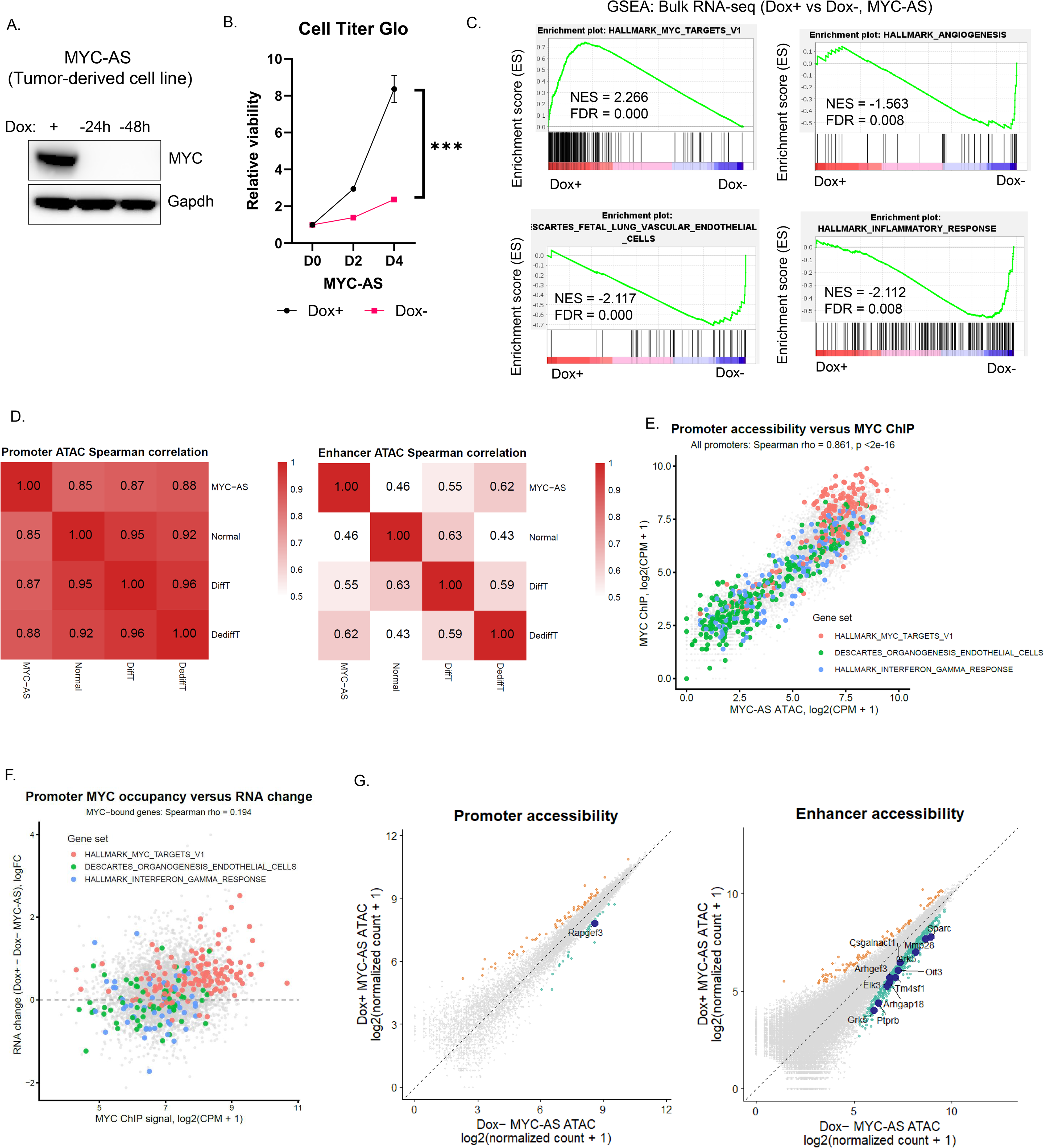
MYC binds accessible promoters and activates canonical growth programs in angiosarcoma. **(A)** Western blot of MYC in MYC-AS cells after 24 or 48h of doxycycline withdrawal. **(B)** Growth curves of *MYC*-active (Dox^+^) and *MYC*-inactive (Dox^-^) MYC-AS cells measured using Cell Titer Glo. Mean ± SD, n=4. *** p < 0.001 at day 4 (Two-tailed unpaired t-test). **(C)** GSEA plots for the indicated gene signatures comparing Dox^+^ and Dox^-^ MYC-AS cells **(D)** Spearman correlations of ATAC-seq signal between MYC-AS cells and *in vivo* Normal, DiffT, and DediffT populations. MYC-AS bulk ATAC-seq signal was quantified across an *in vivo*-defined regulatory atlas generated from merged snATAC-seq peaks identified in the three populations. Correlations were calculated separately for promoter and enhancer regions. **(E)** Scatterplot comparing Dox^+^ MYC-AS ATAC-seq and MYC ChIP-seq signals across *in vivo*-defined promoter regions. Promoters associated with the selected gene sets are highlighted. Spearman correlation coefficient and p-value are shown. **(F)** Scatterplot comparing MYC ChIP-seq signal at MYC-bound promoters with RNA expression changes between Dox^+^ and Dox^-^ MYC-AS cells. Each point represents a gene associated with a MYC-bound promoter. Genes belonging to the indicated gene sets are highlighted. **(G)** Scatterplots comparing promoter (left) and enhancer (right) accessibility between Dox^+^ and Dox^-^ MYC-AS cells. Each point represents one *in vivo*- defined regulatory region. Enlarged dots indicate differentially accessible enhancers associated with endothelial genes (adjusted p < 0.05 and absolute log2 fold change > 0.5; DESeq2). Labeled genes denote endothelial-associated enhancers whose linked genes show concordant RNA upregulation following *MYC* inactivation.

To determine which *in vivo* angiosarcoma chromatin state was closely recapitulated by MYC-AS cells, we performed bulk ATAC-seq on Dox^+^ MYC-AS cells and generated state-specific pseudobulk accessibility peaks using snATAC data from the *in vivo* Normal, DiffT, and DediffT populations. Peak-overlap analysis revealed extensive sharing of promoter peaks, whereas enhancer peaks were more state-restricted (**supp. Fig. 5A**). We next constructed an *in vivo* regulatory atlas from the union of snATAC peaks identified in Normal, DiffT, and DediffT populations and quantified MYC-AS ATAC-seq signal across these regions. At promoters, MYC-AS cells exhibited high correlations with all three *in vivo* cell states, consistent with broad maintenance of promoter accessibility (**Fig. 5D**). At enhancers, there were more peaks from pseudo-bulking of *in vivo* snATAC data than the direct ATAC-seq of MYC-AS *in vitro*, possibly due to technical reasons (**supp. Fig. 5A**). When the enhancer analysis was restricted to regions accessible in MYC-AS cells, MYC-AS accessibility correlated most strongly with DediffT, followed by DiffT and Normal, indicating that the cell line retains a dedifferentiated tumor-associated enhancer landscape.

We next investigated the relationship between MYC occupancy and chromatin accessibility within the angiosarcoma regulatory landscape. Because FACS-sorted *in vivo* tumor populations yielded insufficient material for ChIP-seq, we performed MYC ChIP-seq in Dox^+^ MYC-AS cells. Genome-wide peak annotation showed that MYC binding was enriched at promoter regions (**supp Fig. 5B**). We then quantified MYC ChIP-seq and ATAC-seq signals from Dox^+^ MYC-AS cells across the *vivo*-defined promoter regions. MYC occupancy strongly correlated with promoter accessibility, indicating that MYC preferentially engages accessible promoter elements in MYC-AS cells (**Fig. 5E**). Max, the obligate DNA-binding partner of MYC, co- localized with MYC at these accessible promoter regions, consistent with canonical MYC/Max transcriptional regulation (**supp. Fig. 5C**).

To determine whether promoter-bound MYC directly activates transcription, we integrated MYC occupancy with the gene expression changes following doxycycline withdrawal. Hallmark MYC target genes exhibited high MYC promoter occupancy and were robustly upregulated in Dox^+^ relative to Dox^-^ cells, indicating direct activation of canonical growth programs by MYC in angiosarcoma (**Fig. 5F**). Binding and Expression Target Analysis (BETA) analysis, which integrated MYC ChIP-seq peaks with differentially expressed genes between Dox^+^ and Dox^-^ cells, similarly predicted that MYC functions predominantly as a transcriptional activator in AS cells (**supp. Fig. 5D**) [21].

We next investigated how *MYC* overexpression was associated with repression of endothelial and inflammatory transcriptional programs. BETA analysis did not support widespread direct transcriptional repression by MYC (**supp. Fig. 5D**). We found that genes, exemplified by genes in “Descartes Organogenesis Endothelial Cells” and “Hallmark Interferon Gamma Response” gene sets, generally exhibited lower chromatin accessibility and lower MYC occupancy at their promoters (**Fig. 5E-F**), suggesting that their repression was not mediated by direct promoter binding by MYC. We therefore examined distal regulatory elements that gained accessibility following *MYC* inactivation. Integration with bulk RNA-seq revealed enrichment of endothelial genes linked to focal enhancer regions that became more accessible in Dox^-^ cells and whose expression increased following *MYC* inactivation (**Fig. 5G**). Enhancers preferentially accessible in Dox^+^ cells exhibited greater MYC occupancy, whereas enhancers that gained accessibility in Dox^-^ cells showed comparatively lower MYC binding (**supp. Fig. 5E**). Together, these findings indicate that MYC directly activates canonical growth programs, whereas *MYC* inactivation permits partial restoration of endothelial identity accompanied by focal gains in enhancer accessibility.

### MYC impairs endothelial differentiation in human angiosarcoma

To determine whether the phenotypes observed in the GEMM were recapitulated in human angiosarcoma, we analyzed a transcriptomic dataset comprising tumors with high *MYC* expression (*MYC*-high, n=11) and low *MYC* expression (*MYC*-low, n=11) [22] (**Fig. 6A**). *MYC*- high tumors exhibited increased expression of MYC-associated and proliferative genes (*DKC1*, *CDK4*, *MKI67*) and reduced expression of endothelial genes (*FLT1*, *SLCO2A1*, *VWF*, *CAV1*). Ten of 11 *MYC*-high tumors harbored *MYC* amplification, and all 11 were radiation- or lymphedema-associated angiosarcomas (**Fig. 6B**). GSEA showed enrichment of Hallmark MYC target and cell-cycle and depletion of endothelial lineage signatures in *MYC*-high tumors, consistent with the transcriptional programs observed in the murine models (**Fig. 6C and supp. Fig. 6A and 6B**).

**Figure 6:**
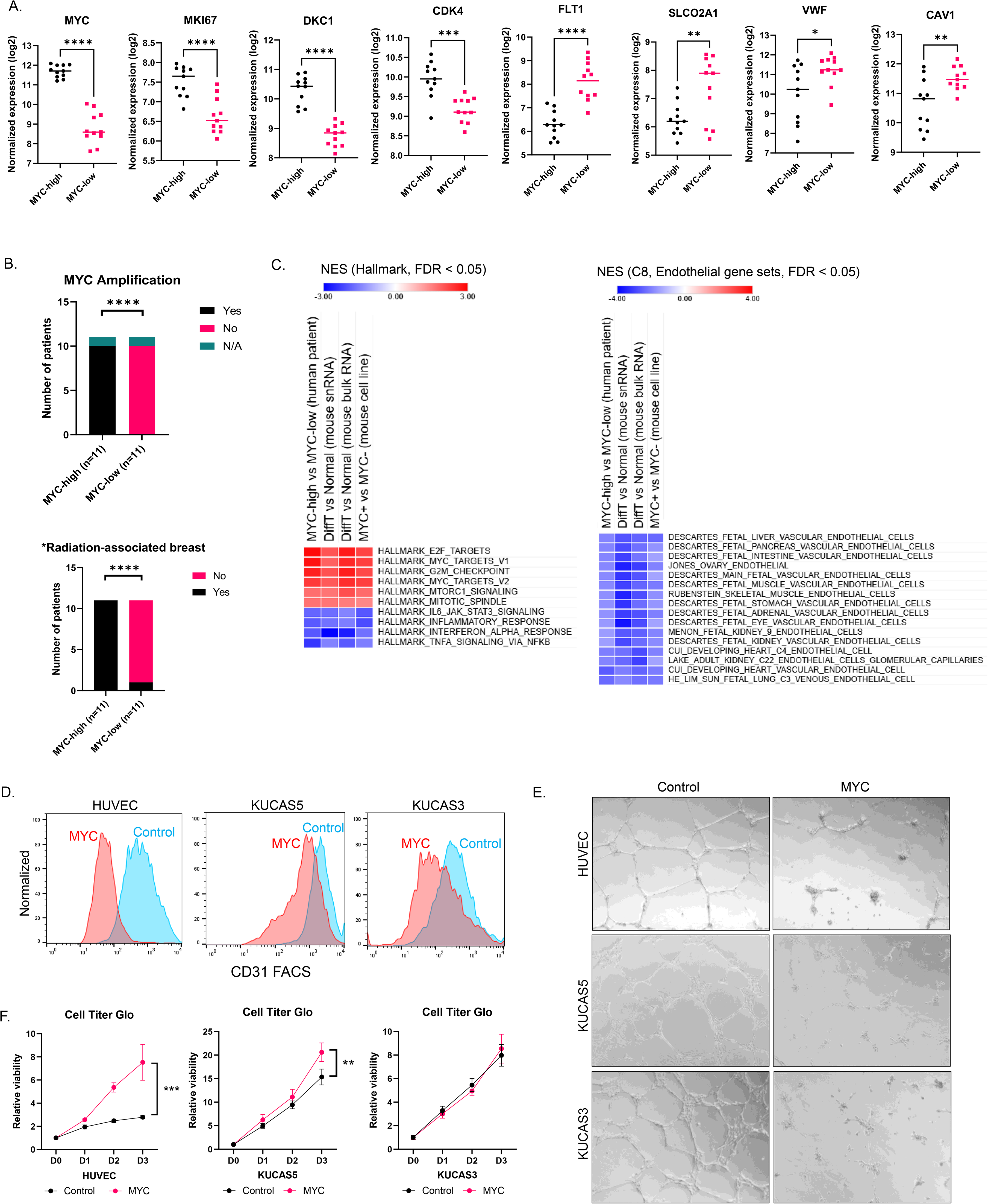
MYC impairs endothelial differentiation in human angiosarcoma. **(A)** Expression of the indicated MYC-associated and endothelial genes in *MYC*-high (n=11) and *MYC*-low (n=11) human angiosarcomas. *p < 0.05, ** p < 0.01, *** p < 0.001, **** p < 0.0001 (Two-tailed unpaired t-test). **(B)** Distribution of *MYC* amplification status and radiation/lymphedema-associated etiology between *MYC*-high and *MYC*-low human angiosarcomas. **** p < 0.0001 (Two-sided Fisher’s exact test). **(C)** Heatmaps showing enrichment scores for Hallmark MYC target, proliferation, endothelial, and interferon-response gene signatures in human angiosarcomas and murine AS cells. **(D)** FACS analysis of cell-surface CD31 expression in HUVEC, KUCAS5, and KUCAS3 cells expressing vector control or MYC- T58A. **(E)** Endothelial tube formation assays of HUVEC, KUCAS5, and KUCAS3 cells expressing vector control or MYC-T58A. **(F)** Growth curves of HUVEC, KUCAS5, and KUCAS3 cells expressing vector control or MYC-T58A. Mean ± SD, n=5. ** p < 0.01, *** p < 0.001 (Two-tailed unpaired t-test).

To test whether MYC directly impairs endothelial differentiation in human cells, we overexpressed a stabilized MYC mutant (MYC-T58A, hereafter MYC) in human umbilical vascular endothelial cells (HUVECs) and two well-differentiated angiosarcoma cell lines, KUCAS5 and KUCAS3 [23] (**supp. Fig. 6C**). FACS analysis showed reduced cell surface expression of the endothelial marker CD31 and the MHC class I component B2M following *MYC* overexpression (**Fig.6D and supp. Fig. 6D**). Transcriptomic analysis of these three models showed enrichment of Hallmark MYC target signatures and depletion of endothelial lineage signatures following *MYC* overexpression (**supp. Fig. 6E**). *MYC* overexpression also reduced endothelial tube formation in all three models (**Fig. 6E**) and increased the proliferation of HUVEC and KUCAS5 cells *in vitro* (**Fig. 6F**). Together, these findings show that *MYC* activation promotes proliferative programs and impairs endothelial identity in human endothelial and angiosarcoma cells.

## DISCUSSION

*MYC* amplification is a defining molecular alteration in most radiation-associated, secondary AS and is also detected in a subset of *de novo* primary tumors [4–6]. Consistent with this association, a molecular subtype enriched for secondary AS exhibited elevated *MYC* expression and oncogenic signaling together with relatively low immune-related signaling [24]. However, studies of MYC as an oncogenic driver in AS have been hampered by the lack of suitable models. Here, our transgenic mouse model demonstrated that *MYC* overexpression alone in adult endothelial cells was sufficient to drive pulmonary AS. The resulting tumors were highly proliferative and progressively lost endothelial lineage and immune-related programs, including endothelial markers and transcription factors that regulate endothelial identity. These findings suggest that MYC promotes transformation not only by driving proliferation but also by disrupting the differentiated endothelial state. A similar loss of lineage identity has been observed in basal-like breast cancer cells following *MYC* overexpression [25] and across MYC-driven tumors arising in the liver, kidney, lung, and spleen [26]. Consistent with our murine findings, *MYC*-high human AS tumors, nearly all of which harbored *MYC* amplification, exhibited increased MYC target and proliferative programs and reduced endothelial lineage signatures. Enforced *MYC* expression in human endothelial and AS cells similarly impaired endothelial differentiation and, in selected models, increased proliferation. Importantly, *MYC* inactivation in murine allografts rapidly suppressed tumor growth and restored endothelial lineage gene expression, demonstrating that both tumor maintenance and the dedifferentiated state remained dependent on sustained MYC activity. Mechanistically, MYC occupancy in MYC-AS cells was strongly associated with promoter accessibility, particularly at promoters accessible across the Normal, DiffT, and DediffT states. Despite widespread MYC occupancy at accessible promoters, MYC-dependent expression was concentrated among canonical MYC target genes with strong promoter occupancy. BETA analysis further predicted that MYC functioned predominantly as a transcriptional activator. These findings are consistent with previous work showing that MYC selectively activates conserved biosynthetic programs that support cell growth [27].

The suppression of inflammatory programs may have important consequences for tumor-immune interactions. MYC has been implicated in tumor immune evasion [28], including reduced MHC-I expression [16–18] and repression of interferon signaling in pancreatic, breast, and lung cancers [29–31]. Similarly, MYC-driven murine AS cells exhibited reduced interferon-response and antigen presentation signatures, accompanied by decreased cell-surface MHC-I expression. Because endothelial cells regulate inflammatory responses and leukocyte trafficking [32], suppression of these programs may alter interactions between AS cells and the tumor microenvironment. Immune-related signatures were also reduced in *MYC*-overexpressing AS cells cultured in the absence of immune and stromal cells, indicating that at least part of this transcriptional response is tumor-cell autonomous. Conversely, *MYC* inactivation restored interferon-response and antigen presentation programs, increased tumor cell MHC-I expression, and was accompanied by changes in intratumoral immune-cell populations and CD8^+^ T-cell states. Nevertheless, these findings do not establish whether the immune changes contribute directly to tumor regression or arise secondarily from MYC-dependent growth arrest and tumor remodeling. Further studies will be required to define the mechanisms through which MYC regulates antigen presentation and to determine how tumor-intrinsic MYC activity influences immune and stromal populations in the AS microenvironment.

## MATERIALS AND METHODS

### Mouse models and tissue dissociation

All mouse studies were approved by the MSKCC Institutional Animal Care and Use Committee (IACUC) under protocol 11-12-029. *VE-Cad*::*CreERT2, CAG-LSL*-*rtTA3-IRES-mKate2*, and *TetO*-*MYC* mice have been previously described [8, 10, 11]. To induce Cre-mediated recombination, 6- to 8-week-old mice received two intraperitoneal injections of tamoxifen (3 mg per dose in corn oil) on days 1 and 3. To activate rtTA3-dependent *MYC* expression, mice were provided drinking water containing 0.5 mg/ml doxycycline and 1% sucrose.

Lungs and subcutaneous allografts were dissected, minced with a scalpel, and digested with 2.5 mg/mL of Collagenase A (Millipore Sigma 10103586001) and 1.25 mg/mL of Dispase II (Millipore Sigma 04942078001) in HBSS supplemented with 20mM HEPES for 1 hour at 37°C with rotation. Cell suspensions were filtered through a 70-μm cell strainer and treated with 1x red blood cell lysis buffer (Biolegend 420301).

### Flow cytometry and fluorescence-activated cell sorting

Single cell suspensions from murine tissues were blocked with TruStain FcX anti-mouse CD16/32 antiobody (BioLegend Clone 93) and stained with the indicated combinations of APC- Cy7 anti-mouse CD45 (BioLegend Clone 30-F11), Alexa Fluor 488 anti-mouse CD31 (BioLegend Clone 390), PE anti-mouse H-2 (MHC-I) (BioLegend Clone M1/42), Alexa Fluor 488 anti-mouse CD8a (BioLegend Clone 53-6.7), and PE anti-mouse CD4 (BioLegend GK1.5) DAPI or 7-AAD was used to exclude dead cells. Human cells were blocked with Human TruStain FcX (BioLegend 422302) and stained with APC/Cyanine7 anti-human CD31 (Biolegend Clone WM59) or APC/Cyanine7 anti-human B2M (Biolegend Clone A17082A). Flow cytometry and cell sorting were performed using BD LSRFortessa, BD FACSAria III, or BD FACSymphony S6 as appropriate.

### Histology, immunohistochemistry, and multiplex immunofluorescence

Mouse tissues were fixed overnight with 4% paraformaldehyde at 4°C, paraffin embedded, and sectioned. Sections were stained with hematoxylin and eosin or subjected to immunohistochemistry using a Ventana Discovery XT platform. Primary antibodies used for immunohistochemistry were rabbit anti-MYC (Abcam ab32072, 1:500), rabbit anti-Ki67 (Abcam ab16667, 1:100), rabbit anti-mKATE (Origene TA150072, 1:4000), rabbit anti-Erg (Abcam ab92513, 1:1000), rabbit anti-CD8a (Cell Signaling Technology 98941, 1:100), and rabbit anti-CD4 (Abcam ab183685, 1:1000). Multiplex immunofluorescence was performed using a Leica Biosystems Bond RX platform with rabbit anti-MYC (Abcam, ab32072, 0.125 µg/ml), rabbit anti-mKATE (Origene TA150072, 1:250), and rabbit anti-Erg (Abcam ab92513, 0.125 µg/ml).

### Western blotting

Cells cultured *in vitro* were lysed in RIPA buffer (Pierce 89901) supplemented with 1x Halt Protease and Phosphatase Inhibitor Cocktail (Thermo Scientific 1861281). Protein lysates were combined with LDS sample buffer and DTT, heated at 75°C, resolved on NuPAGE 4-12% gels, and transferred to PVDF membranes. Membranes were blocked and incubated with primary antibodies overnight at 4°C. Primary antibodies were rabbit anti-MYC (Abcam ab32072, 1:1000), rabbit anti-MAX (Abcam ab199489, 1:1000), and mouse anti-GAPDH (ProteinTech 60004-1-Ig, 1:3000).

### Single-nucleus Multiome sequencing

Equals number of live (7-AAD^-^) murine endothelial cells (CD45^-^/CD31^+^/mKATE2^-^), differentiated AS cells (CD45^-^/CD31^+^/mKATE2^+^), and dedifferentiated AS cells (CD45^-^/CD31^-^/mKATE2^+^) were isolated from enzymatically dissociated lung tissues by FACS and pooled. Fresh nuclei were extracted and processed using the 10x Genomics Chromium Next GEM Single Cell Multiome ATAC + Gene Expression platform according to the manufacturer’s protocol. Sequencing reads were mapped using the “cellranger-arc” software (10x Genomics) to a custom mouse genome (mm10) containing the *mKATE_Bglobin*, *MYC_SV40pA*, and *CreER^T2^* transgenes.

For RNA analysis, cell-containing droplets were identified using scCB2 [33], and putative doublets were removed using the DoubletDetection package. Nuclei with fewer than 1,000 or more than 40,000 total RNA counts, greater than 20% mitochondrial reads, or fewer than 200 detected genes were excluded. Genes detected in fewer than 20 nuclei and all mitochondrial genes were removed. Principal component analysis (PCA), UMAP dimensionality reduction, and Leiden clustering were performed using Scanpy as previously described. Cell-type annotation was performed using CellTypist with Tabula Muris as the reference dataset [14, 15]. For each cell state, the fraction of nuclei assigned to each predicted cell type was calculated.

ATAC-seq data were processed using ArchR [34]. Putative doublets and nuclei with a TSS enrichment score below 10 were removed. The retained nuclei exhibited similar TSS enrichment profiles across the 3 biological samples, with a median TSS enrichment score of 13.3965 and a median of 5,980 fragments per nucleus. A total of 4,866 nuclei with high-quality RNA and ATAC profiles were retained. Iterative latent semantic indexing, UMAP dimensionality reduction, and graph-based clustering of ATAC profiles were performed using ArchR. RNA-defined cell-state annotations were assigned to the corresponding ATAC profiles using shared Multiome cell barcodes and visualized on the ATAC UMAP. ATAC fragments from nuclei assigned to each cell state were aggregated to generate state-specific pseudobulk profiles, and peaks were called separately for the Normal, DiffT, and DediffT populations using ArchR.

Pseudotime trajectory analysis was performed in ArchR using Normal cells as the root state. Gene-associated chromatin accessibility was quantified using the GeneScoreMatrix. Integrated RNA expression was obtained from the GeneIntegrationMatrix, and transcription factor motif accessibility was quantified using motif deviation z-scores from the MotifMatrix. Transcription factor motif enrichment across cell states was calculated using ArchR. Adjusted motif enrichment p values were transformed as −log10(adjusted p value). Motifs with an adjusted p value below 0.05 in at least one cell state were retained, and selected significant motifs were visualized using Morpheus (Broad Institute). A relative color scheme was applied independently to each motif, with the minimum and maximum values across cell states mapped to the endpoints of the color scale.

### Bulk RNA-sequencing

Live (DAPI^-^) murine endothelial cells (CD45^-^/CD31^+^/mKATE2^-^), differentiated AS cells (CD45^-^/CD31^+^/mKATE2^+^), and dedifferentiated AS cells (CD45^-^/CD31^-^/mKATE2^+^) were isolated from enzymatically dissociated lung tissues by FACS. RNA was extracted using TRIzol (Invitrogen, 15596026) and subjected to Ultra-Low Input RNA-sequencing. For cell lines experiments, cells were lysed directly in TRIzol, and the extracted RNA was used for RNA-sequencing. Libraries were prepared using standard Illumina PolyA selection protocol and sequenced on an Illumina HiSeq2500 platform using 50-bp paired-end reads, with 30-40 million reads generated per sample. Sequencing reads were aligned to the mouse (mm10) or human reference genome (hg38) using STAR [35]. Expression values were normalized to total mapped reads. Gene set enrichment analysis was performed using GSEA v4.1.0 program with the Hallmark, C2 curated, and C8 cell type signature gene sets from the Molecular Signatures Database, as well as manually curated endothelial gene sets.

### Cell culture

Murine AS cells (CD45^-^/CD31^-^/mKATE2^+^) were isolated by FACS and transplanted subcutaneously into NOD-SCID mice maintained on doxycycline-containing drinking water. The resulting allografts were dissociated, and tumor cells were cultured on fibronectin (Millipore Sigma F1141)-coated plates in primary mouse endothelial cell medium comprising DMEM: Ham’s F-12 supplemented with 20% FBS, 2 mM GlutaMAX, 20 mM HEPES, 100 μg/mL heparin, nonessential amino acids, penicillin/streptomycin, 5 μM SB431542 (Selleckchem), 10 ng/ml Igf-1 (Millipore Sigma I8779), 10 ng/ml Fgf-2 (Gibco PMG0035), 10 ng/ml Egf (PeproTech AF-100-15), and 10 ng/ml Vegf Millipore (Sigma V4512). Doxycycline was added to the culture medium at 10 ng/ml to maintain *MYC* epxression. The resulting murine cell line, was designated MYC-AS.

HUVECs, provided by Tomer Itkin and Shahin Rafii, and the human angiosarcoma cell lines KUCAS5 (ABM T8029) and KUCAS3 (ABM T8028) were cultured in endothelial cell growth medium (PromoCell C-22010). HUVECs were cultured on fibronectin-coated plates, whereas KUCAS5 and KUCAS3 cells were cultured on gelatin (Stem Cell Technologies 07903)-coated plates.

### Allograft experiments and doxycycline withdrawal

MYC-AS cells (1.0 × 10^6^) were resuspended in 100 μL of 1:1 mix of growth medium and Matrigel (Corning 356237) and injected subcutaneously into 6- to 8-week-old C57BL/6J mice (The Jackson Laboratory) maintained on doxycycline-containing drinking water. After tumors reached approximately 100 mm^3^, mice were randomized to continued doxycycline treatment (Dox^+^) or doxycycline withdrawal (Dox^-^). Tumor length (L), width (W), and height (H) were measured by caliper at the indicated time points, and tumor volume was calculated using the following formula: V = π*(L × W × H)/6. Tumors were collected for histologic, flow cytometric, and single-cell RNA-sequencing analyses.

### Single-cell RNA-sequencing

Live cells (DAPI^-^) were isolated by FACS from enzymatically dissociated allografts. Because mKATE2^+^ tumor cells were scarce in allografts collected 6 days after doxycycline withdrawal, cells from this time point were sorted into mKATE2^+^ tumor cell and mKATE2^-^ non-tumor cell fractions. Equal numbers of cells from the two fractions were combined at a 1:1 ratio before library preparation. The mKATE2^-^ fraction was collected without additional enrichment for specific immune or stromal populations. Single-cell RNA-seq libraries were generated using the 10x Genomics Chromium Next GEM Single Cell 3’ Gene Expression following the manufacturer’s protocol. For each sample, 200 million reads were acquired on an Illumina NovaSeq platform S4 flow cell.

Single-cell RNA-seq data were analyzed as previously described [36, 37]. Sequencing reads were mapped using the Cell Ranger multi software (10X Genomics) to a custom mouse genome (mm10) containing the *mKATE_Bglobin*, *MYC_SV40pA*, and *CreER^T2^* transgenes. Cell- containing droplets were identified using scCB2, and putative doublets were removed using the DoubletDetection package [33]. The levels of mitochondrial reads and numbers of unique molecular identifiers (UMIs) were similar among the samples, indicating that there were no systematic biases in the libraries from different samples. Cells with fewer than 100 detected genes, fewer than 2,000 or more than 50,000 total RNA counts, or greater than 20% mitochondrial reads were excluded. Genes detected in fewer than 20 cells and all mitochondrial genes were removed. Combining samples in the entire cohort of Dox^-^ 0d (n=3), Dox^-^ 2d (n=2), Dox^-^ 4d (n=2) and Dox^-^ 6d (n=3) yielded a filtered count matrix of 57,851 cells by 21,439 genes, with a median of 12,127 counts and a median of 3,172 genes per cell.

Downstream analysis was performed using Scanpy (version 1.9.0) [38]. Counts were normalized by log2(10K+1). The count matrix was further scaled to mean 0 and standard deviation 1 for principal component analysis (PCA) and uniform manifold approximation projection (UMAP) dimension reduction (https://arxiv.org/abs/1802.03426), and Leiden clustering [39]. PCA was performed on the 2,000 most variable genes, and the top 50 principal components (PCs) retained 40.5% of the variance. Cell types were assigned using established marker genes identified from the literature and reference information from PanglaoDB [40]. The mean expression of each cluster and each sample were generated with Scanpy. GSEA was performed using JAVA GSEA v4.1.0 program with the Hallmark, C2 curated, and C8 cell type signature gene sets from the Molecular Signatures Database, as well as manually curated endothelial gene sets. Gene expression imputation for visualization was performed using MAGIC (Markov affinity-based graph imputation of cells) [41].

### Bulk ATAC-sequencing

MYC-AS cells maintained in doxycycline-containing medium or collected 24 hours after doxycycline withdrawal were subjected to bulk ATAC-seq using a standard protocol. Libraries were sequenced on an Illumina platform, and sequencing reads were aligned to the mouse reference genome (mm10) using Bowtie 2 (version 2.4.5) [42]. Accessible chromatin peaks were called using MACS2 [43]. Peaks shared between biological replicates were retained to generate a consensus peak set for each condition. Raw read counts were quantified using featureCounts (v2.0.1) [44], and differential chromatin accessibility between Dox^+^ and Dox^-^ cells was analyzed using DESeq2 [45]. Regions with an adjusted p value < 0.05 and an absolute log2 fold change > 0.5 were considered differentially accessible. and normalized by reads per genome content (RPGC). Positive log2 fold changes indicated greater accessibility in Dox^+^/MYC-on cells, whereas negative values indicated greater accessibility in Dox^-^/MYC-off cells. Genome-normalized signal tracks were generated using reads per genomic content (RPGC), and heatmaps were generated using deepTools v3.5.1 [46].

### ChIP-sequencing

MYC-AS cells maintained in doxycycline-containing medium or collected 24 hours after doxycycline withdrawal were subjected to chromatin immunoprecipitation as previously described [47]. MYC-bound chromatin was immunoprecipitated using rabbit anti-MYC antibody (Cell Signaling Technology 18583), and Max-bound chromatin was immunoprecipitated using mouse anti-Max antibody (Santa Cruz Biotechnology sc-8011). Libraries were generated and sequenced on an Illumina HiSeq4000 platform using 50 bp paired-end reads to a depth of 30-40 million reads per sample. Reads were processed for adapter trimming and aligned to the mouse reference genome (mm10) using Bowtie2. Mapping quality was analyzed with Qualimap (2.2.2- dev) [48]. ChIP-seq peaks were called using MACS2 [43].

### Integrative analysis of ATAC-seq, ChIP-seq, and RNA-seq

ATAC-seq and ChIP-seq peaks were annotated to the nearest TSS using HOMER annotatePeaks [49]. Regions with peak centers located from 1,000 bp upstream to 500 bp downstream of the nearest TSS were classified as promoters; all other regions were classified as enhancers. An *in vivo* regulatory atlas was generated by merging snATAC-seq peaks identified in the Normal, DiffT, and DediffT populations. Normalized Dox+ MYC-AS and *in vivo* ATAC-seq signals were quantified across these regions. Spearman correlations between MYC-AS and the three *in vivo* cell states were calculated separately for promoter and enhancer regions. Genomic overlaps among the MYC-AS and *in vivo* ATAC-seq peak sets were also calculated separately for promoters and enhancers and visualized using UpSet plots.

At promoter regions, MYC-AS ATAC-seq signal was integrated with MYC ChIP-seq signal to assess the relationship between chromatin accessibility and MYC occupancy. MYC and Max ChIP-seq signals were subsequently visualized at promoter regions accessible across the Normal, DiffT, and DediffT states. Promoters were assigned to their nearest genes, and MYC occupancy was integrated with RNA-expression changes between Dox^+^ and Dox^-^ MYC-AS cells to evaluate the relationship between promoter binding and MYC-dependent transcription. Binding and Expression Target Analysis (BETA) was performed by integrating MYC ChIP-seq peaks from Dox+ MYC-AS cells with differential gene-expression data comparing Dox^+^ and Dox^-^ MYC-AS cells [21]. BETA was used to assess whether MYC binding was associated with transcriptional activation or repression and to identify candidate direct MYC target genes.

Differentially accessible enhancers between Dox^+^ and Dox^-^ MYC-AS cells were assigned to the nearest gene, and the linked genes were intersected with endothelial gene sets. Enhancers gaining accessibility in Dox^-^ cells were integrated with RNA-seq data comparing Dox^-^ and Dox^+^ MYC- AS cells to identify linked endothelial genes that also increased in expression following MYC inactivation. MYC ChIP-seq signal was compared between enhancers preferentially accessible in Dox+ cells and those that gained accessibility following doxycycline withdrawal.

### Analysis of human angiosarcoma transcriptomic data

Normalized microarray expression data and associated clinical annotations for human angiosarcoma were obtained from the published dataset [22]. When multiple probes mapped to the same gene, normalized probe-level expression values were averaged to obtain a single gene-level value. Samples were then ranked by gene-level *MYC* expression and divided at the cohort median into *MYC*-high and *MYC*-low groups, resulting in 11 tumors per group.

### Stable MYC Expression

MYC-T58A cDNA was amplified from MSCV Myc T58A puro (Addgene 18773) and cloned into the EcoRI and XhoI sites of MSCV-IRES-GFP (Addgene 20672). Retroviruses encoding MYC-T58A or vector control were produced in 293T cells using standard methods. Viral supernatants were collected at 48- and 72-hours post-transfection, pooled, and filtered through a 0.45 µm PVDF membrane. HUVEC, KUCAS5, and KUCAS3 cells were exposed to viral supernatant in the presence of 8 µg/ml polybrene for 24 hours. GFP-positive cells were isolated by FACS to establish stable populations.

### Cell growth and endothelial tube formation assays

Equal numbers of cells expressing vector control or MYC-T58A were seeded in clear-bottom 96- well plates, with seeding density optimized for each cell line. Cell viability was measured at the indicated timepoints using Cell Titer Glo. For endothelial tube formation assays, equal numbers of cells expressing vector control or MYC-T58A were seeded onto Matrigel (Corning 356231)- coated 24-well plates, and brightfield images were acquired 24 hours later.

### Statistics

All statistical analyses were performed using GraphPad Prism 10 (GraphPad Software) unless otherwise indicated. Comparisons between two independent groups were performed using two-tailed unpaired t-tests with Welch’s correction. Comparisons among three or more groups were performed using one-way ANOVA followed by Tukey’s multiple-comparisons test. Categorical variables were compared using two-sided Fisher’s exact tests; samples with unavailable data were excluded from the corresponding analysis. Statistical tests, sample sizes, and definitions of error bars are specified in the figure legends. Adjusted p values were used where indicated for multiple-testing analyses. A p value below 0.05 was considered statistically significant.

## Data availability

Sequencing data generated in this study will be deposited in the Gene Expression Omnibus. Accession information will be provided upon availability.

## Acknowledgements

We thank the Chi and Chen labs for critical discussion and feedback. We also thank the Flow Cytometry Core Facility, Molecular Cytology Core Facility, and Integrated Genomics Operation Core Facility for facilitating various experiments.

---

**supp. Fig. 1:**
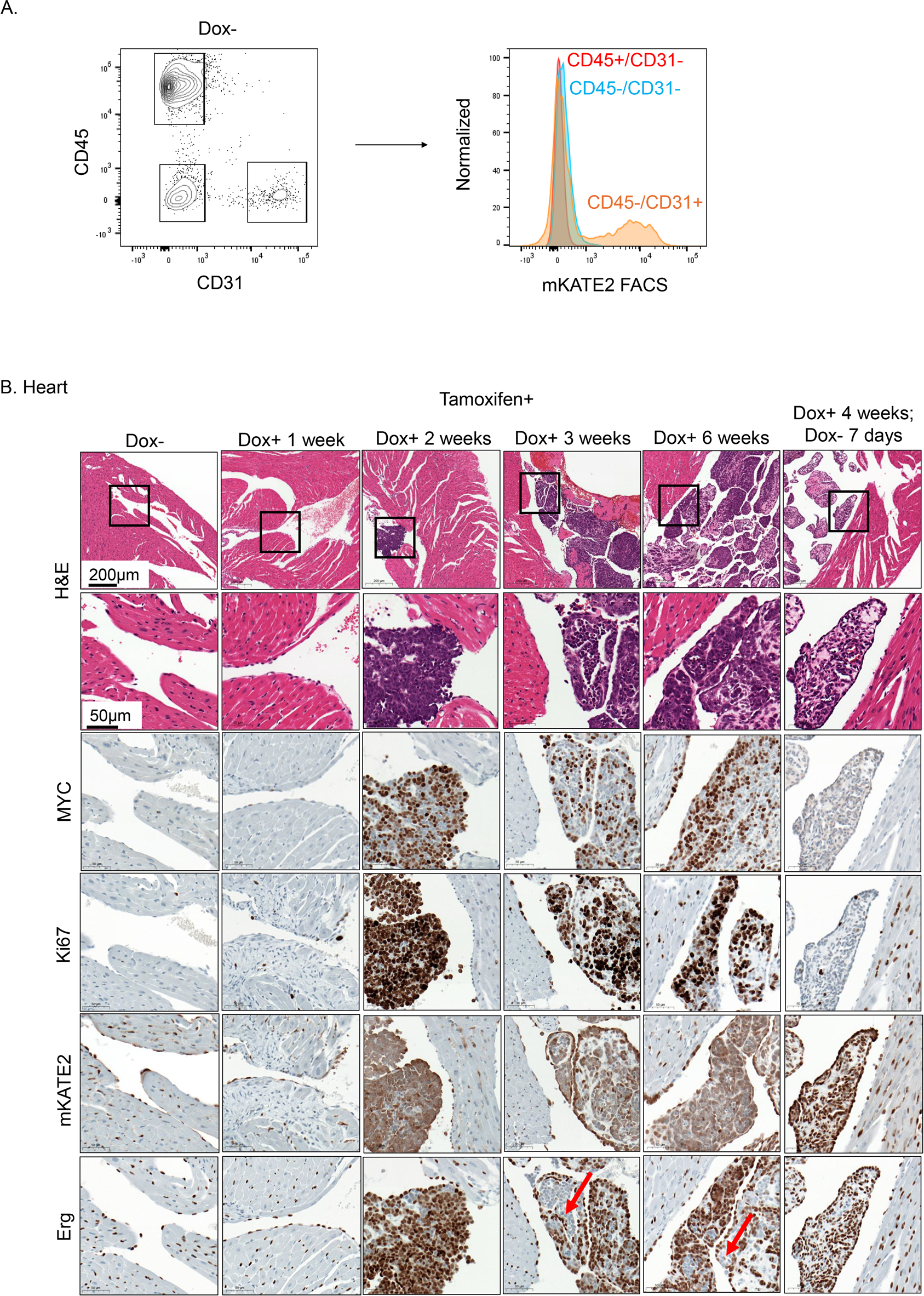
**(A)** Histogram of mKATE2 fluorescence in 1) CD45^+^/CD31^-^, 2) CD45^-^/CD31^-^, and 3) CD45^-^/CD31^+^ populations from dissociated lung cells of a representative tamoxifen-treated, doxycycline-naive mouse. **(B)** H&E and IHC staining of heart tissues from tamoxifen-treated mice after the indicated durations of doxycycline administration. The arrow indicates Erg^-^ tumor cells.

**Supp. Fig. 2:**
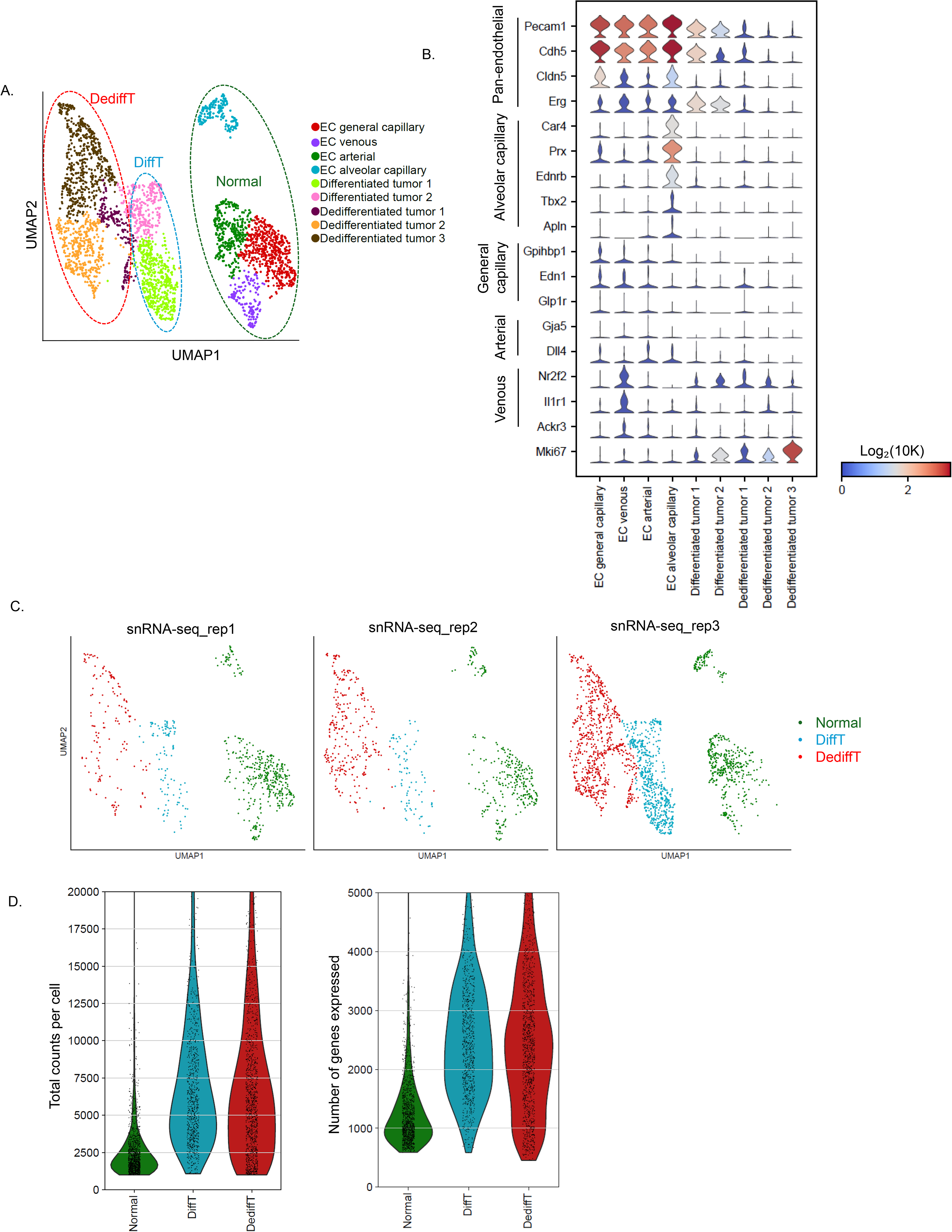
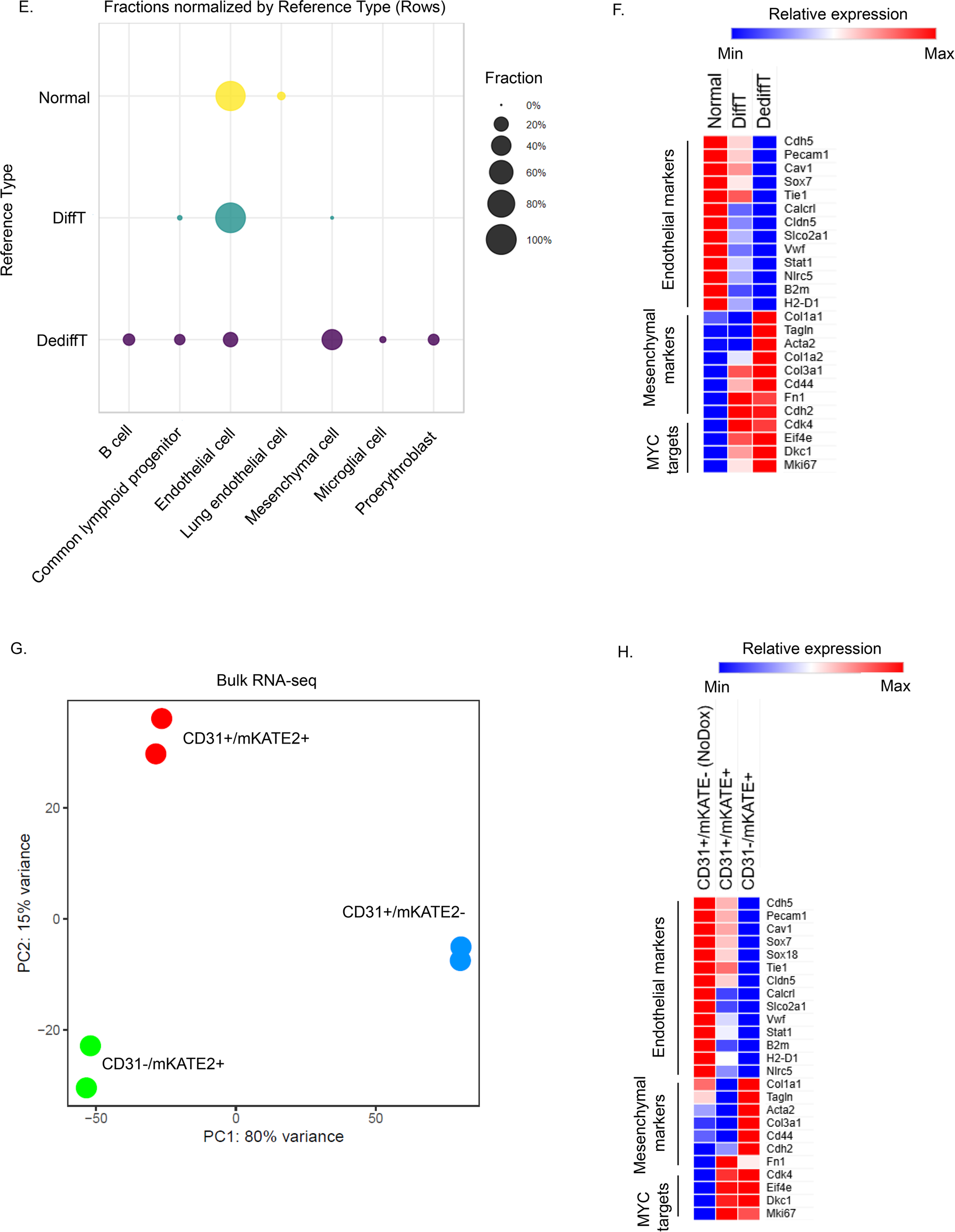
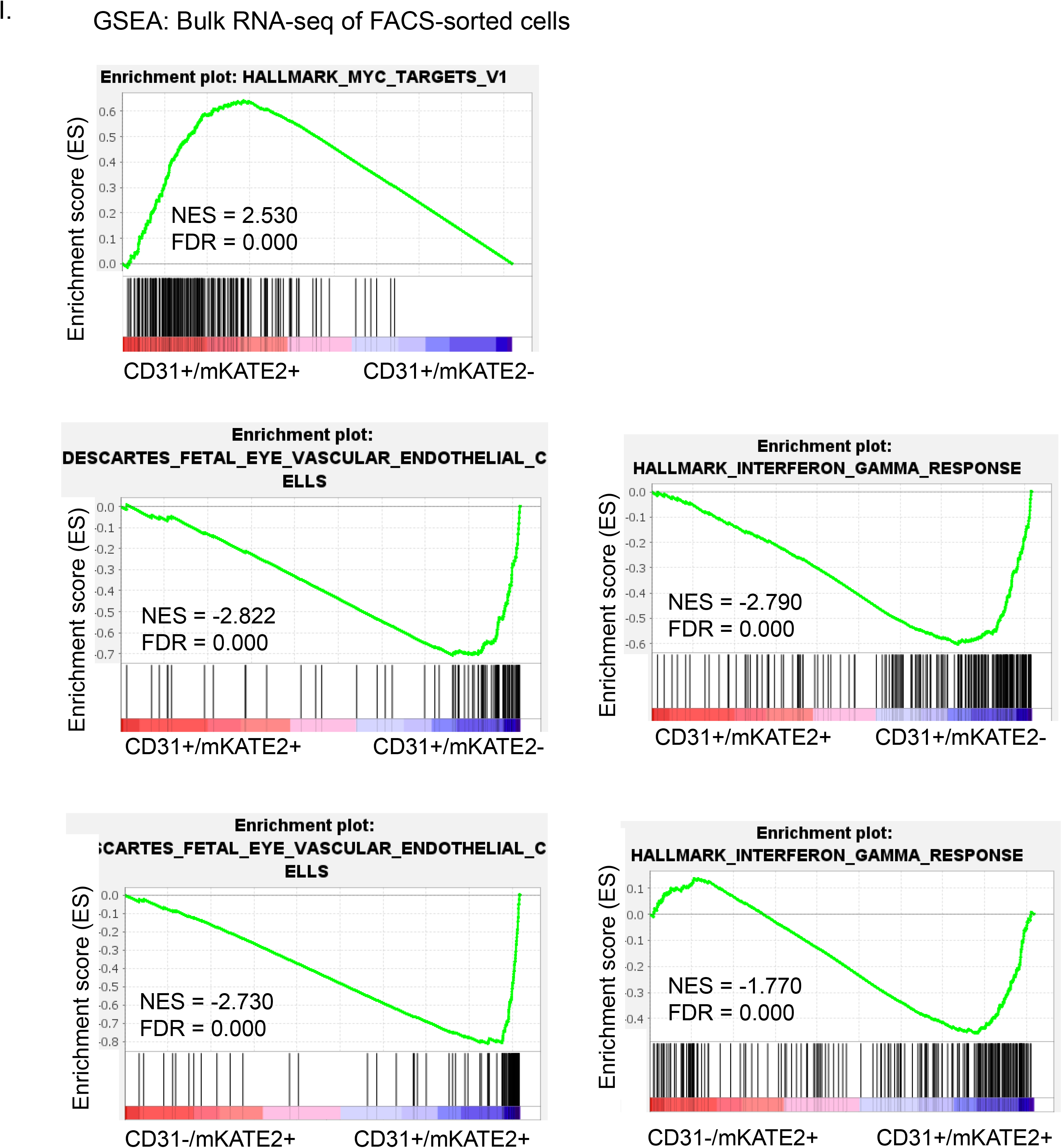
**(A)** snRNA-seq UMAP showing the eight clusters identified by Leiden clustering. Cluster grouping used to define Normal, DiffT, and DediffT states are indicated. **(B)** Violin plots showing normalized expression of the indicated endothelial subtype-specific genes across the eight clusters. **(C)** snRNA-seq UMAPs showing the distribution of Normal, DiffT, and DediffT nuclei separately for each mouse. **(D)** Violin plots showing total RNA counts and number of detected genes per nucleus in the Normal, DiffT, and DediffT states. **(E)** Dot plot showing the fractions of nuclei in each cell state assigned to the indicated cell types by CellTypist using Tabula Muris as the reference dataset. Dot size represents the fraction of nuclei assigned to each predicted cell type. **(F)** Heatmap showing relative expression of selected endothelial, mesenchymal, and MYC target genes across the three cell states. **(G)** Principal component analysis of bulk RNA-seq profiles from FACS-sorted CD31^+^/mKATE2^-^, CD31^+^/mKATE2^+^, and CD31^-^/mKATE2^+^ populations. **(H)** Heatmap showing relative expression of endothelial, mesenchymal, and MYC target genes in the three FACS-sorted populations. **(I)** GSEA plots for the indicated gene signatures comparing CD31^+^/mKATE2^+^ and CD31^-^/mKATE2^+^ tumor cells with CD31^+^/mKATE2^-^ normal endothelial cells.

**Supp. Fig. 3:**
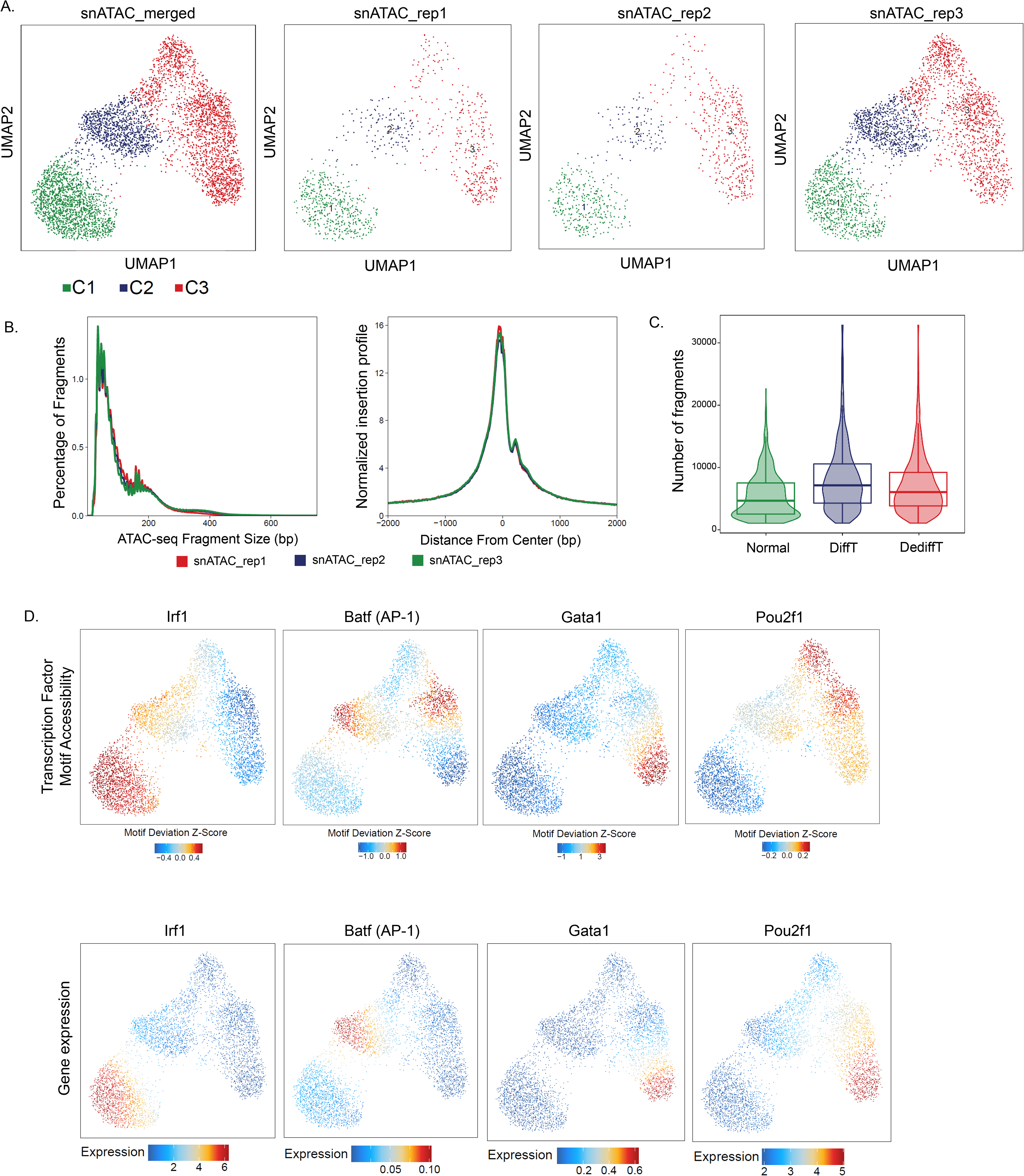
**(A)** snATAC-seq UMAP showing the distribution of cells collected from each mouse (n=3) across the three major clusters. **(B)** ATAC-seq fragment-size distributions (left) and transcription start site enrichment profiles (right) for the three biological replicates. **(C)** Violin plots showing the number of unique ATAC-seq fragments per cell in the Normal, DiffT, and DediffT cell states. **(D)** UMAPs showing transcription factor motif accessibility (top; chromVAR motif deviation z-score) and integrated gene expression of the corresponding transcription factors (bottom; GeneIntegrationMatrix normalized counts).

**Supp. Fig. 4:**
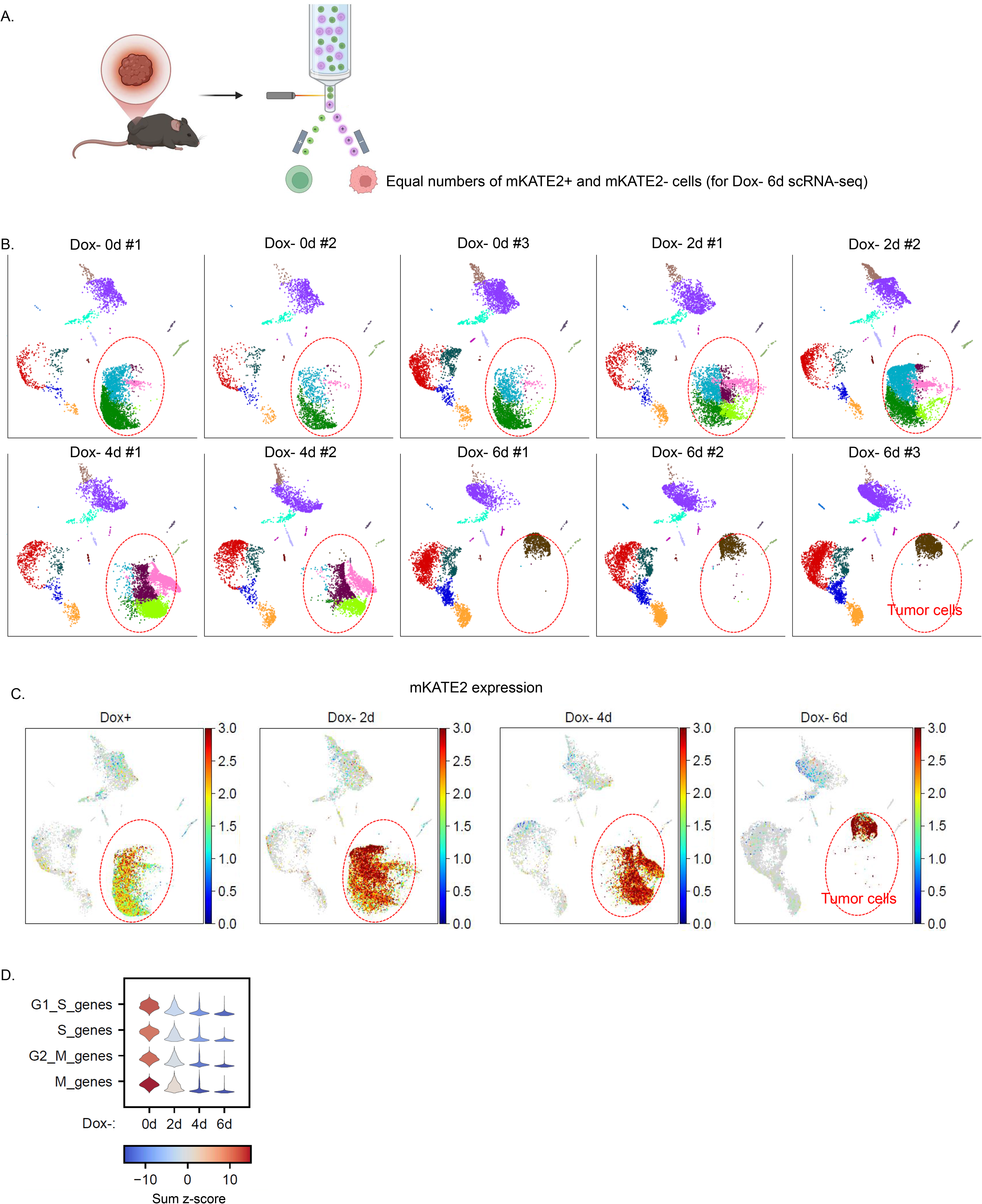
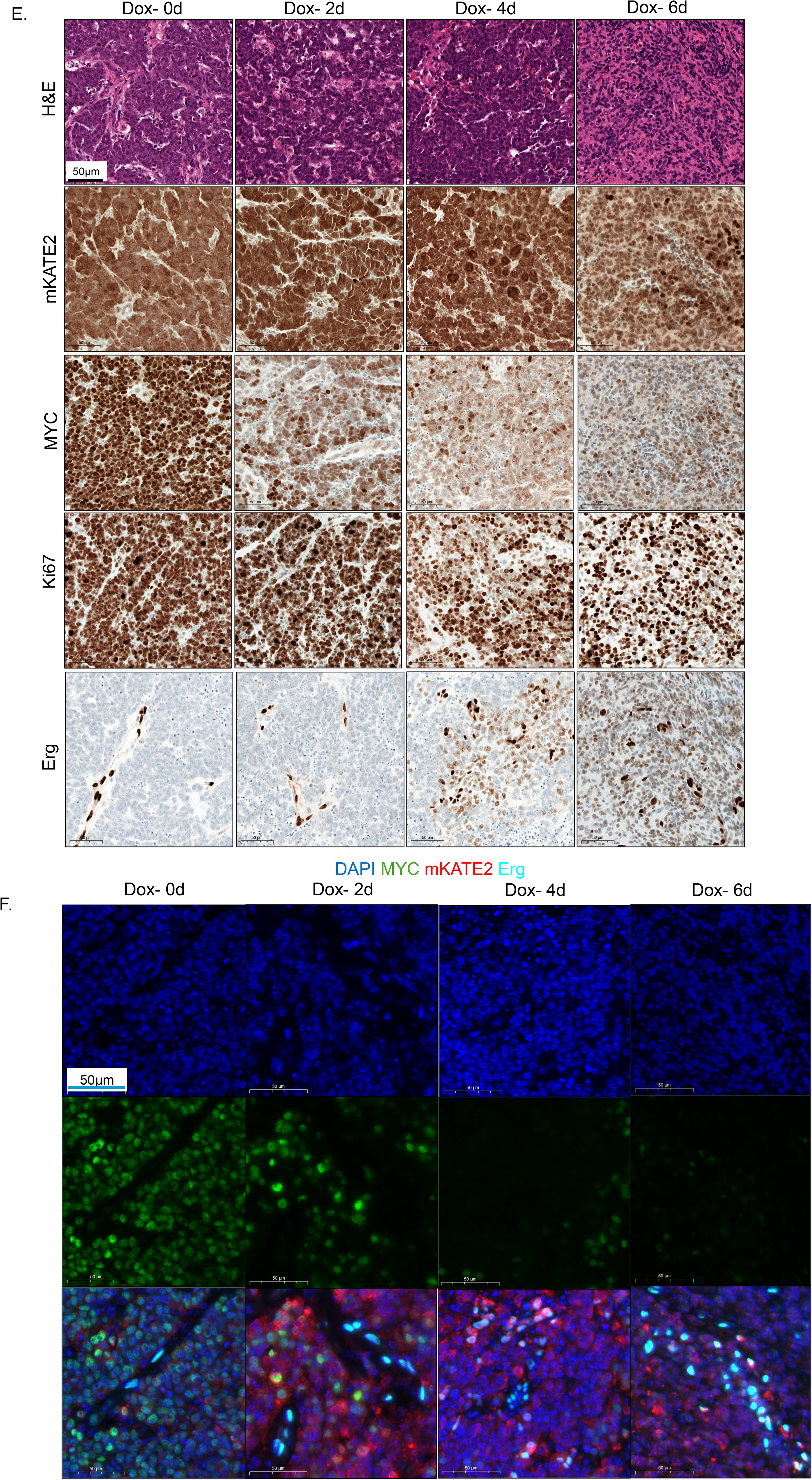
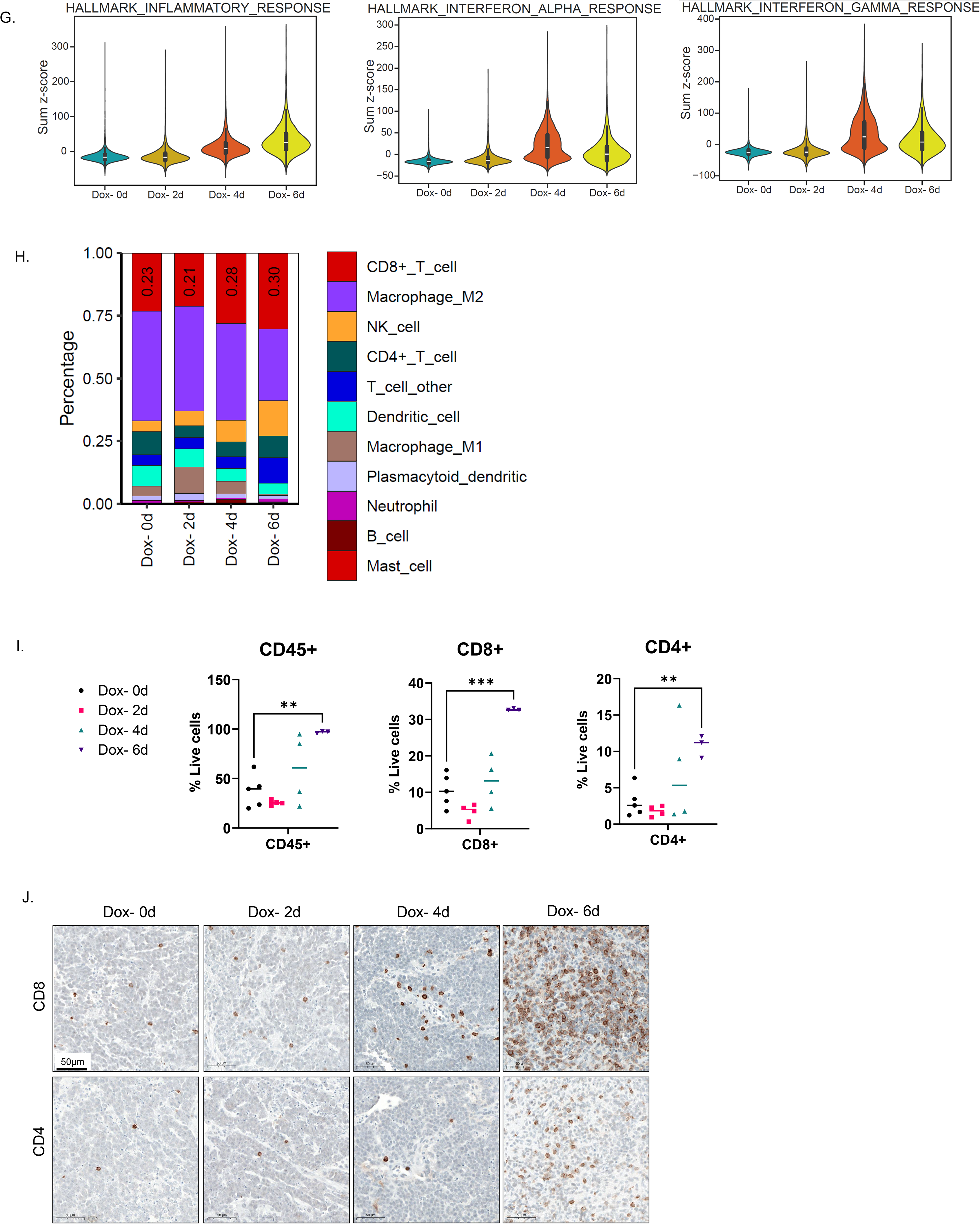
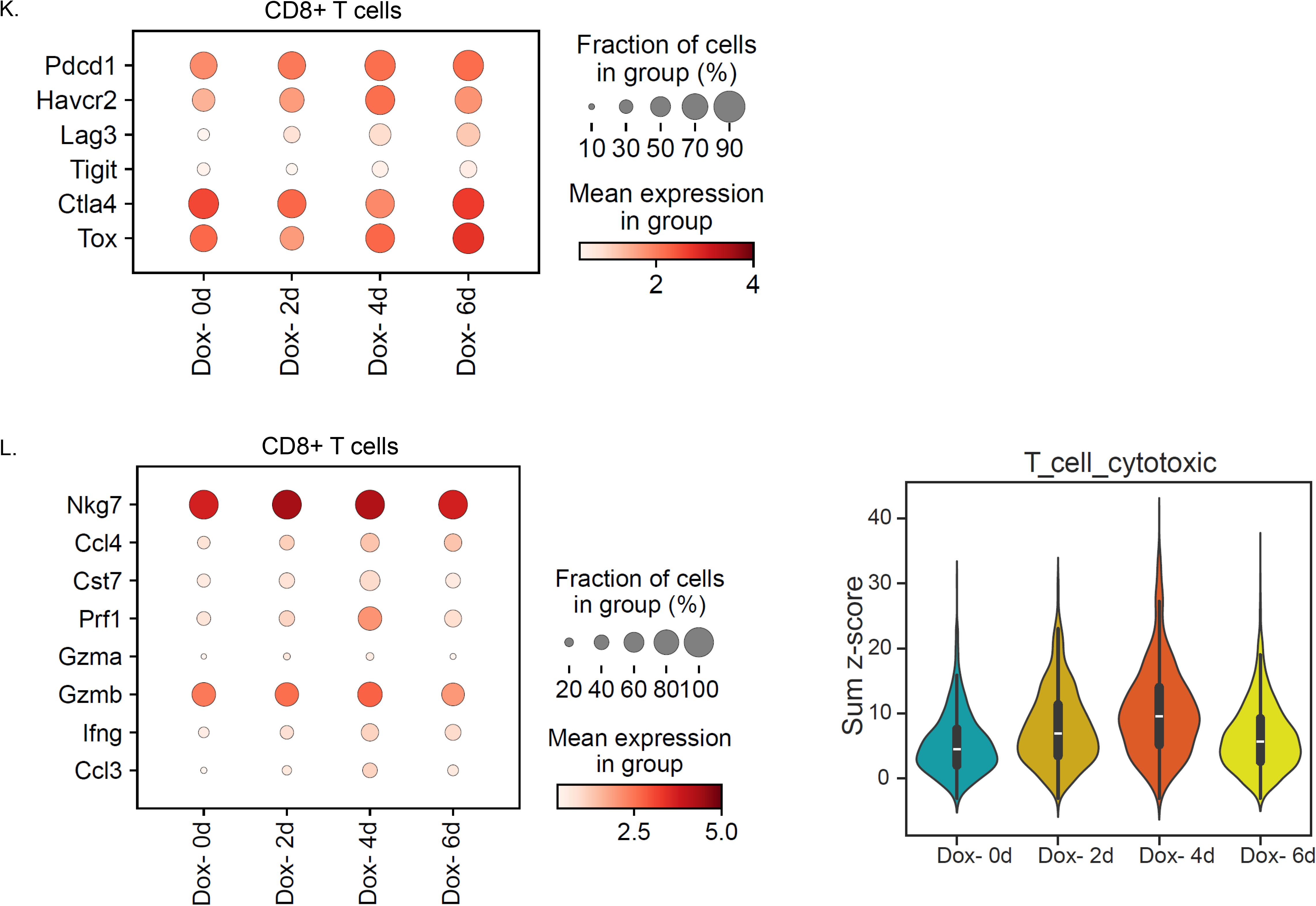
**(A)** FACS-isolation strategy. For allografts after 6 days of doxycycline withdrawal, equal numbers of mKATE2^+^ tumor cells and mKATE2^-^ non-tumor cells were isolated and pooled before scRNA-seq. **(B)** scRNA-seq UMAPs shown separately for each allograft sample and colored by cell-cluster identity. **(C)** scRNA UMAPs colored by *mKATE2* expression. **(D)** Sum z-scores for cell-cycle genes in tumor cells at the indicated times after doxycycline withdrawal. **(E)** Immunohistochemical staining of the indicated proteins in allografts at the specified times after doxycycline withdrawal. **(F)** Multiplex immunofluorescence staining of the indicated proteins in AS allografted collected at the indicated times after doxycycline withdrawal. **(G)** Violin plots showing sum z-scores for the indicated immune-related programs in the tumor cells at the specified times after doxycycline withdrawal. **(H)** Relative abundance of the indicated immune cell populations within the non-tumor compartment at the specified times after doxycycline withdrawal. **(I)** Flow cytometry quantification of CD45^+^, CD8^+^, and CD4^+^ cells among live (DAPI^-^) cells in allografts at the indicated times after doxycycline withdrawal. **p < 0.01, ***p < 0.001 (Day 0 vs day 6, two-tailed unpaired t-test). **(J)** Immunohistochemical staining of the indicated proteins in AS allografts at the indicated times after doxycycline withdrawal. **(K)** Dot plot showing the expression of the indicated exhaustion-associated markers in intratumoral CD8^+^ T cells at the indicated times after doxycycline withdrawal. **(L)** Dox plot showing expression of indicated cytotoxic effector markers (left) and violin plot showing their sum z-scores (right) in intratumoral CD8^+^ T cells at the specified times after doxycycline withdrawal

**Supp. Fig. 5:**
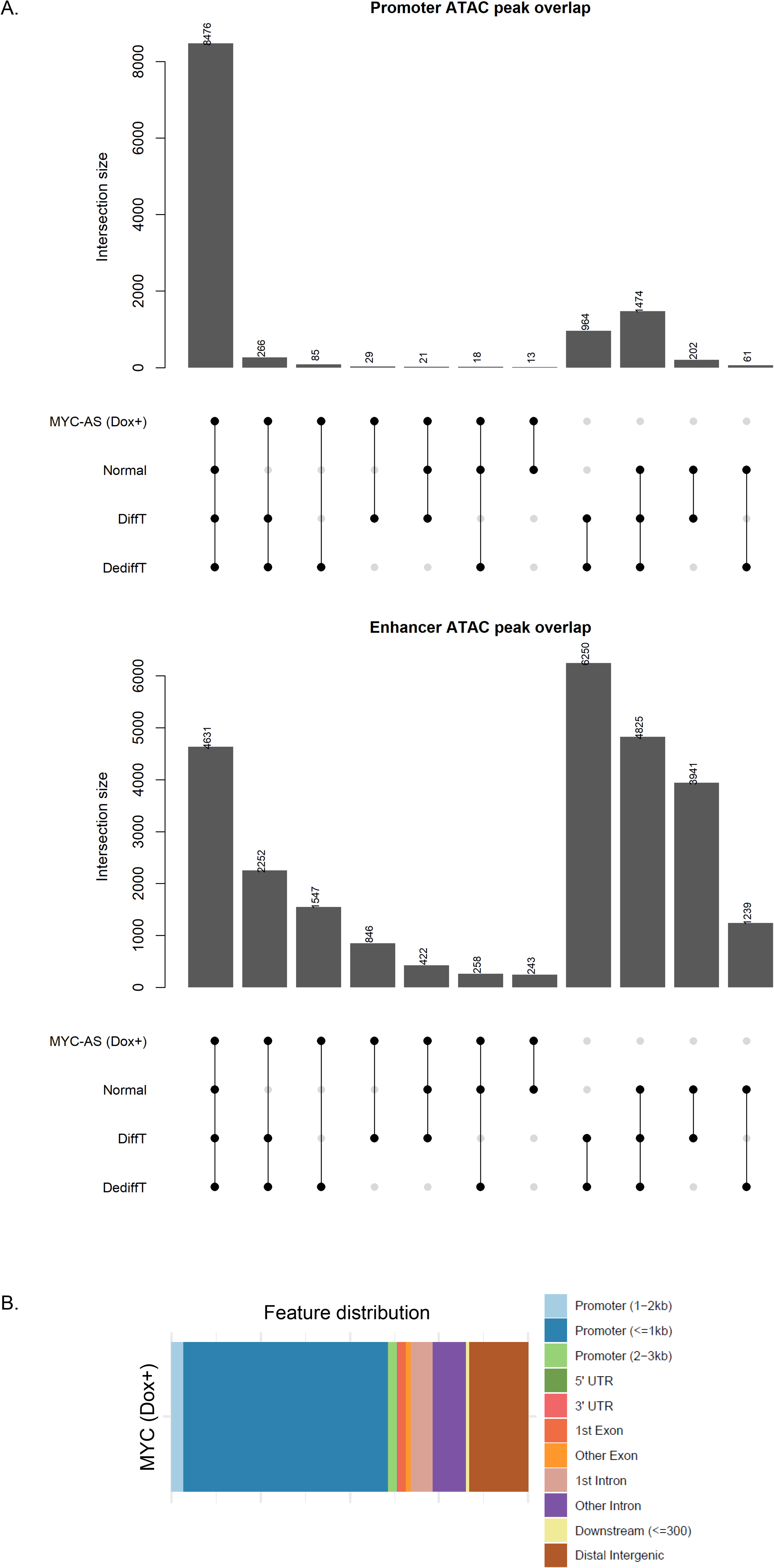
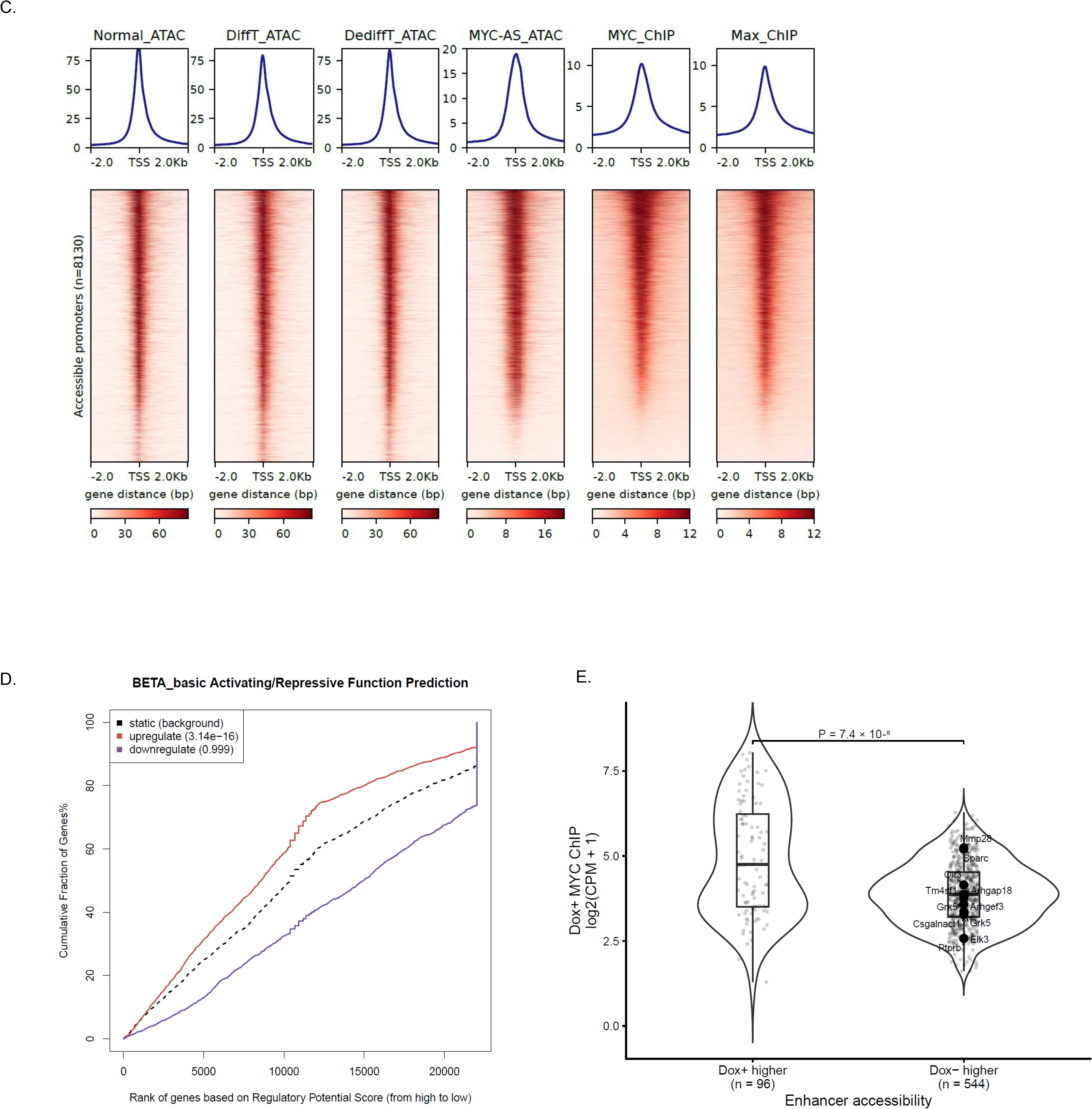
**(A)** UpSet plots showing overlap among promoter and enhancer ATAC-seq peaks identified in MYC-AS cells and the *in vivo* Normal, DiffT, and DediffT populations. **(B)** Genomic annotation of MYC ChIP-seq peaks in MYC-AS cells. **(C)** Heatmaps of MYC and Max ChIP-seq signals at promoter regions accessible across the *in-vivo* cell states. **(D)** Binding and Expression Target Analysis (BETA) integrating MYC ChIP-seq peaks with differentially expressed genes between Dox^+^ and Dox^-^ MYC-AS cells. **(E)** Violin plots comparing MYC ChIP-seq signal between enhancers preferentially accessible in Dox^+^ cells (n=96) and enhancers that gained accessibility following doxycycline withdrawal (Dox^-^, n=544). Each point represents one enhancer. Labeled points indicate enhancers linked to the selected endothelial genes.

**Supp. Fig. 6:**
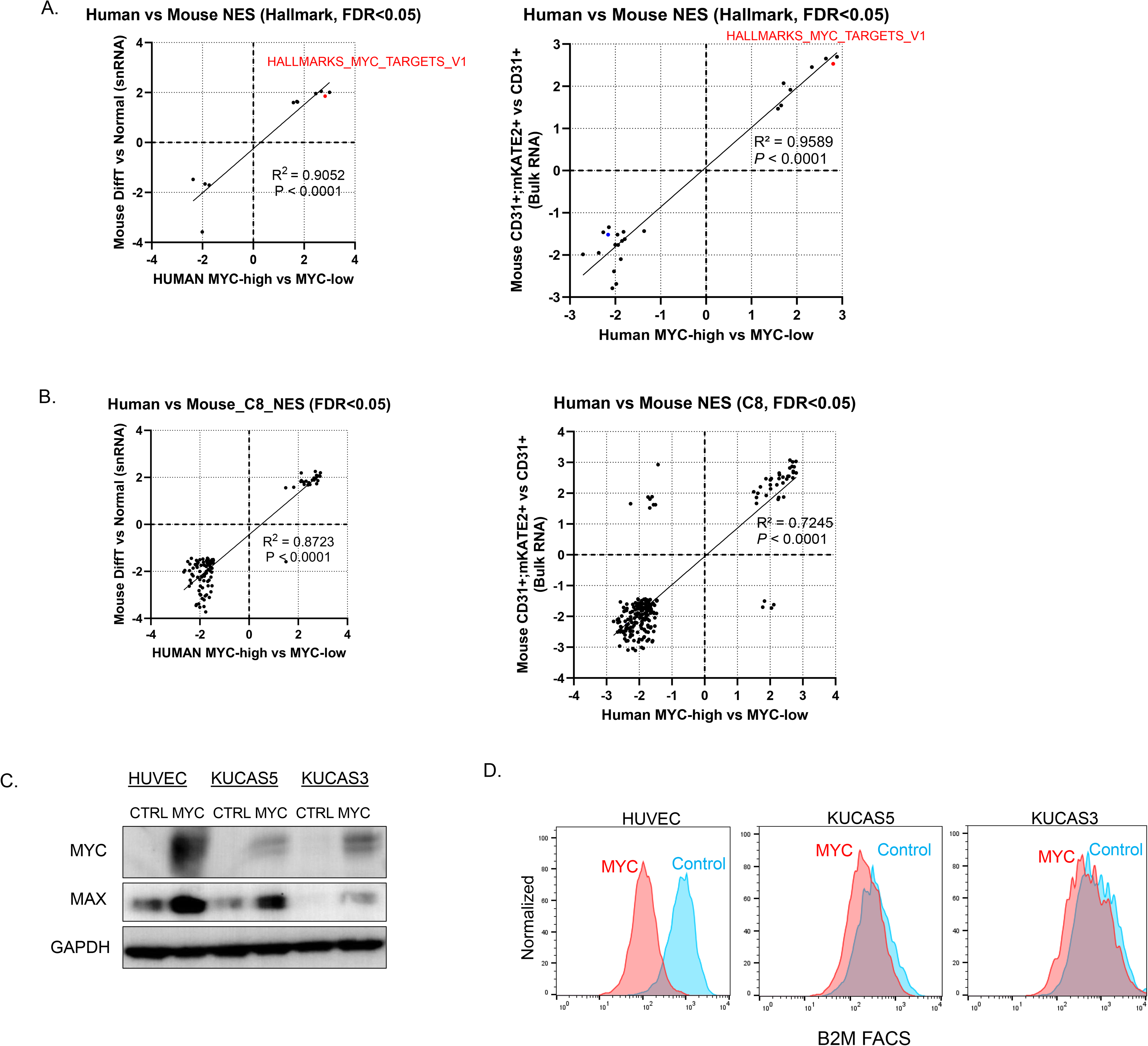
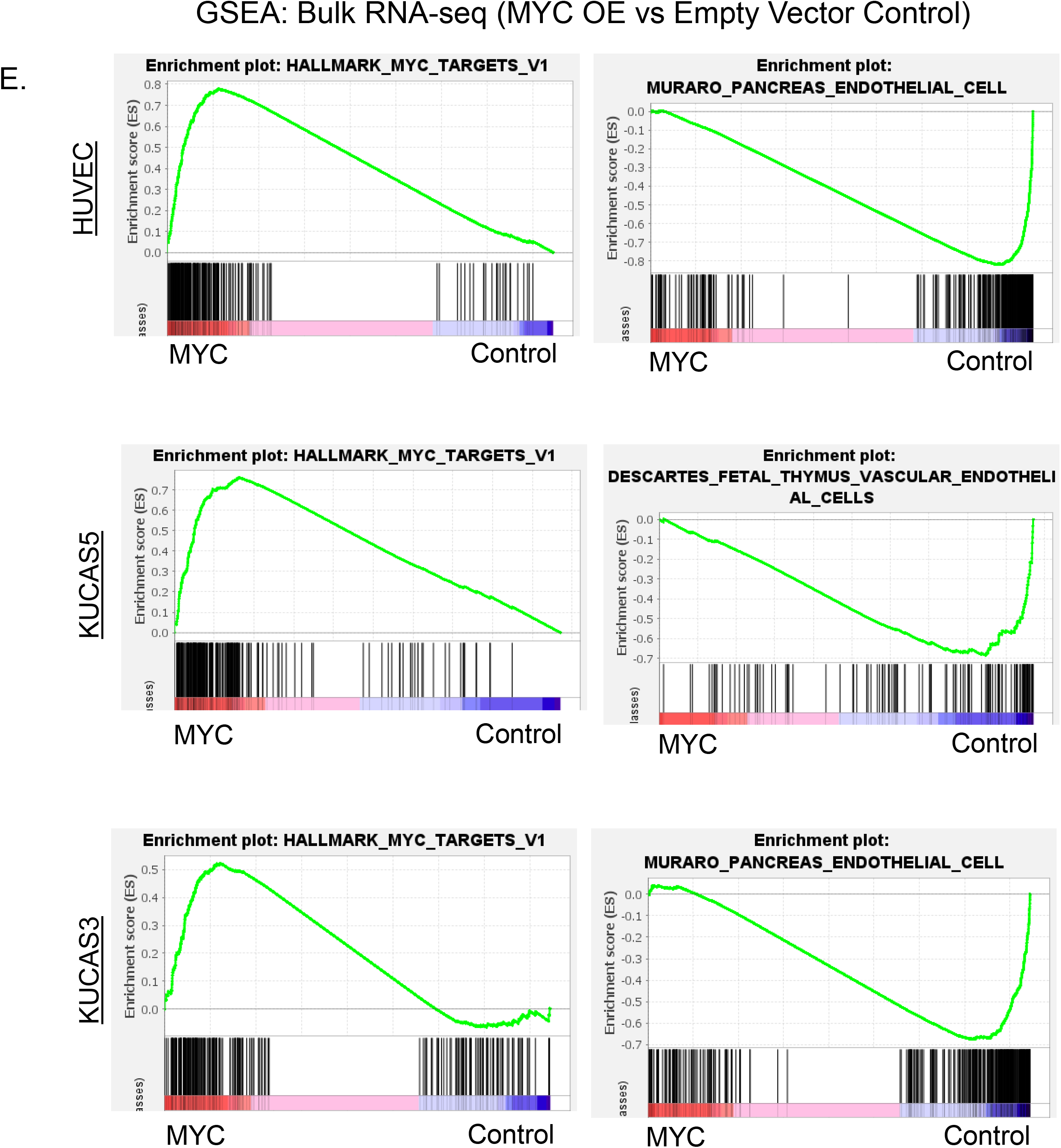
**(A)** Correlation of Hallmark gene sets enrichment scores between *MYC*-high human angiosarcomas and murine AS tumors. Each point represents one gene set. The Spearman correlation coefficient and p value are shown. **(B)** Correlation of C8 cell-type signature enrichment scores between *MYC*-high human angiosarcomas and murine AS tumors. Each point represents one gene set. The Spearman correlation coefficient and p value are shown. **(C)** Western blot analysis of MYC in HUVEC, KUCAS5, and KUCAS3 cells expressing vector control or MYC-T58A. **(D)** FACS analysis of cell-surface B2M expression in HUVEC, KUCAS5, and KUCAS3 cells expressing vector control or MYC-T58A. **(E)** GSEA plots for the indicated Hallmark MYC target and endothelial lineage signatures in HUVEC, KUCAS5, and KUCAS3 cells expressing vector control or MYC-T58A.

